# Extrinsic Noise or Intrinsic Coupling: Dissecting Correlated Fluctuations in Gene Transcription

**DOI:** 10.1101/2025.03.17.643613

**Authors:** Shihe Zhang, Huihan Bao, Heng Xu

## Abstract

Understanding stochastic gene transcription requires distinguishing between intrinsic molecular randomness and extrinsic environmental variability. Traditionally, these components are identified as the independent and correlated fluctuations between identical gene copies within the same cell. However, intrinsic gene-gene interactions can introduce additional correlations, challenging this standard approach. Here, we develop a new noise decomposition framework based on a theoretical model of stochastic transcription for an intrinsically coupled gene pair under fluctuating environments. By analytically deriving correlated fluctuations of nascent RNA, we disentangle intrinsic coupling from extrinsic noise based on their distinct relationships with the mean and variance of gene activation probabilities across varying environments. Applying this framework to single-cell transcription data from *Drosophila* embryos, we uncover previously unidentified couplings between sister alleles and between alternative promoters of an endogenous gene. Our findings offer a versatile approach for deciphering intrinsic gene-gene interactions in stochastic transcription within fluctuating environments.

## Introduction

Gene transcription, the production of RNA in biological cells, is a well-known stochastic process driven by both the intrinsic randomness of biomolecular reactions and extrinsic environmental variability [1–4]. Understanding the mechanisms of stochastic transcription kinetics requires accurately distinguishing these two components within the total fluctuation (or noise). Formally, this is done by decomposing the variance of cellular RNA number *m* using the law of total variance [4–10]:

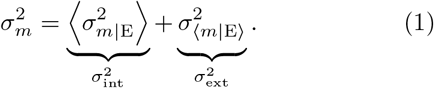

Here, the intrinsic variance 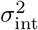 measures the average fluctuation in *m* under fixed environmental conditions (E), whereas the extrinsic variance 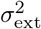 captures the variability in the conditional average of *m* due to fluctuating environments.

To experimentally achieve such partitioning, the standard approach is the dual reporter assay, which monitors two distinguishable copies of the same gene embedded in the same cell (via, e.g., fluorescent RNA reporter) [1,5,6,9,11–13]. Assuming each gene copy transcribes independently, the covariance between their RNA numbers reflects extrinsic variance from the shared environment, while the remaining uncorrelated variance is intrinsic.

However, while this simple approach has been the gold standard for two decades, the assumption of gene copy independence appears overly idealistic, as genes within the same cell inevitably interact. For instance, gene copies may compete for transcription factors [14,15], and their products may exert feedback regulations [16, 17]. Moreover, homologous alleles of the same gene located on different chromosomes within the same cell can cross-regulate each other via direct trans-homolog enhancer-promoter interactions, a phenomenon known as transvection [18,19]. Similar enhancer-sharing mechanisms also occur between genes on the same chromosome or between promoters of the same gene [20,21]. In principle, these interactions can introduce couplings between genes, even in the absence of extrinsic fluctuations, thereby violating the independence assumption.

In many biological systems, particularly in eukaryotic cells where genes typically exist in homologous pairs, these effects may be nonnegligible [13,21]. While some indirect effects, such as feedback regulations, can be separated using short-lived reporters (e.g., nascent RNA reporters) or time-correlation analysis [13,22], most direct couplings at the gene activation level are inseparable from the system. Attempting to eliminate or ignore these intrinsic couplings is often both technically cumbersome and scientifically counterproductive, particularly when studying endogenous genes in realistic (rather than over-simplified synthetic) systems. Therefore, a new framework is needed to disentangle correlated transcriptional fluctuations arising from intrinsic gene-gene coupling and extrinsic noise. The goal of this Letter is to develop such a framework.

### The model

We start with a simple telegraph model for nascent transcription of a pair of intrinsically coupled identical genes in a eukaryotic cell [23–26] [Fig. 1(a), see Supplemental Material [27]]. In this model, each gene copy (*i* = *a, b*) fluctuates between an active (ON) and an inactive (OFF) transcription state, with baseline transition rates *k*_ON_ and *k*_OFF_. If one gene is ON, it modulates the activation rate of its sister copy by a factor *γ*, i.e., *k*_ON_ *→ γk*_ON_. Here, *γ* = 1 indicates independent gene copies, while *γ >* 1 and 0 *< γ <* 1 denote positive and negative couplings, respectively. When a gene is active, it initiates new transcription events as a Poisson process with rate *k*_INI_, followed by a constant-speed (*V*_EL_) RNA synthesis (of length *L*) and a post-synthesis residence duration *T*_S_ before the nascent RNA is released.

**FIG. 1.**
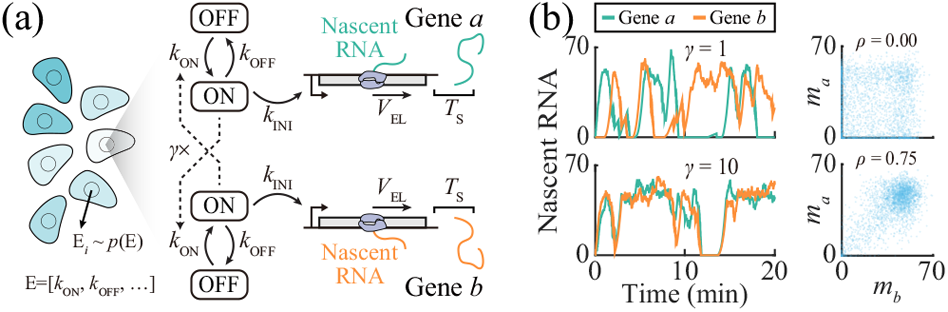
A stochastic model of coupled gene transcription. (a) Model schematic. (b) Simulated results for the model with and without intrinsic coupling under a fixed environmental condition. Left: Time traces of gene transcription. Right: Scatter plot of nascent RNA levels for both gene copies. *γ* ≠ 1 leads to correlated fluctuation. The kinetic parameters are set as: *k*_ON_ = 0.5 min^−1^, *k*_OFF_ = 0.5 min−1, *k*_INI_ = 50 min^−1^, *T*_RES_ = 2 min, *g*(*τ*) = −*τ*, and *γ* = 1 and 10 for independent and coupled gene pairs, respectively.

To account for extrinsic variability, we assume that all kinetic parameters E = [*k*_ON_, *k*_OFF_, …] may vary across cells and over time following a probability distribution *p*(E) [Fig. 1(a), see Supplemental Material [27]]. Based on prior studies, many of these extrinsic variations occur on longer time scales than nascent RNA dynamics, justifying an adiabatic approximation, with quasi-static E over the duration of nascent RNA production [49,50]. Among the parameters in E, *k*_ON_ and *k*_OFF_ are likely the primary targets of extrinsic fluctuations [4,50]. Simulations of this model demonstrate that two gene copies with *γ ≠* 1 can generate correlated outputs even in the absence of extrinsic fluctuation [Fig. 1(b)].

### Solving coupled intrinsic fluctuation

At a given time *t*, the state of the system is characterized by four random variables: the gene states *n*_*i*_ (0 and 1 for OFF and ON states, respectively) and the amount of nascent RNA *m*_*i*_ (*m*_*i*_ ≥ 0) at each gene copy [23, 51]. Notably, *m*_*i*_ represents the cumulative signal (in units of a complete RNA molecule) from all nascent RNA molecules initiated within the residence time window *T*_RES_ = *L/V*_EL_ + *T*_S_ prior to *t*, i.e., 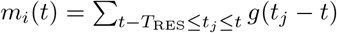 [23] (Supplemental Material [27]). Here, *g*(*τ*) represents the contribution of a nascent RNA molecule initiated at time *τ* (relative to *t*, − *T*_RES_ ≤ *τ* ≤ 0) to the total nascent RNA signal at *t*. Note that the system can be nondimensionalized by rescaling *τ* and all kinetic rates with respect to *T*_RES_. Without loss of generality, we set *T*_RES_ = 1 for this study. The probability distribution of the system state 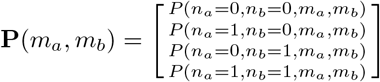 and its statistical moments can be obtained analytically (Appendix A) [23,52,53].

Specifically, under a fixed environmental condition, the steady-state marginal distribution of *n*_*a*_ and *n*_*b*_ is:

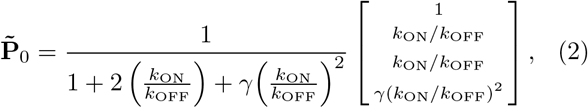

which yields the covariance between *n*_*a*_ and *n*_*b*_ [Fig. 2(a)]:

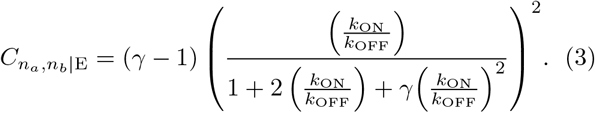

**FIG. 2.**
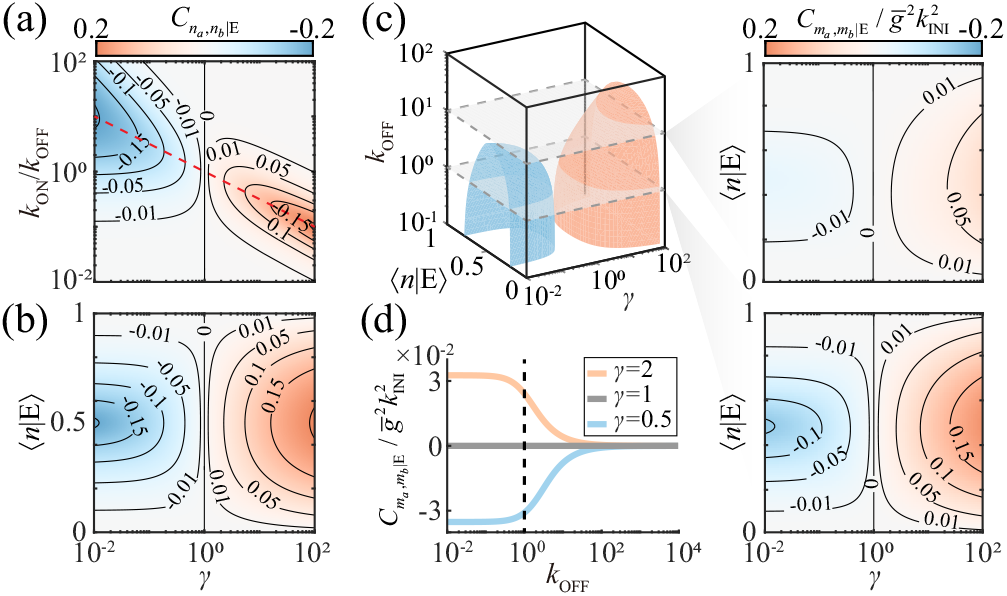
Intrinsic covariance in coupled gene transcription. (a–b) Covariance of gene state as a function of *γ* and *k*_ON_*/k*_OFF_ (a) or *γ* and ⟨*n*|E⟩ (b) under a fixed environmental condition. (c) Covariance of nascent RNA as a function of *γ*, ⟨*n*|E⟩, and *k*OFF. Red and blue surfaces, 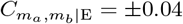, respectively. (d) Decay of nascent-RNA covariance with *k*_OFF_ for representative *γ* values (⟨*n*|E⟩ = 0.5).

As expected, *γ >* 1 and 0 *< γ <* 1 lead to positive and negative covariance, respectively. For each *γ* ≠1, Eq. (3) reveals an extremum of 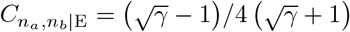, corresponding to equal probabilities of the ON and OFF states for a single gene copy, i.e., ⟨*n*|E⟩ = 0.5. Equivalently, 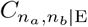 can be expressed as a function of ⟨*n*|E⟩ and *γ* [Fig. 2(b)], i.e., 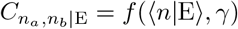, with (Supplemental Material [27]):

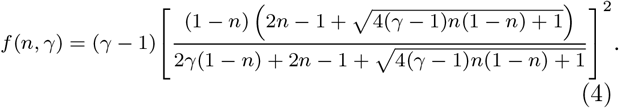

Because ⟨*n*|E⟩ is experimentally more accessible than *k*_ON_ and *k*_OFF_, this reformulation helps connect our theory to experimental data in subsequent derivations. In addition to covariance, the total variance of *n* is simply 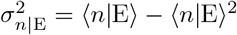, independent of *γ*.

These gene-state quantities are closely related to experimentally measurable nascent RNA statistics (Appendix A). For example, the mean nascent RNA signal is proportional to the ON-state probability:

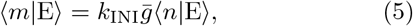

where 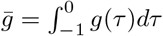. Furthermore, both the covariance and total variance at nascent RNA level follow trends similar to those at the gene-state level [Fig. 2(c) and S1(a)]. Specifically, in the limit of slow gene-state transitions (*k*_ON_ & *k*_OFF_ → 0), the covariances and total variance at these two levels are simply related as:

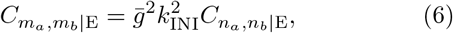

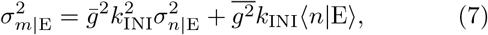

where 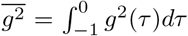. Notably, these approximations hold well up to *k*_ON_ & *k*_OFF_ ∼ 1 [Fig. 2(d) and S1(b), see Appendix A]. Therefore, for experiments measuring short-lived nascent RNA signals, whose lifetimes satisfy *k*_ON_ & *k*_OFF_ ≲ 1 in many biological systems [54], Eqs. (6) and (7) remain valid.

### Partitioning correlated noise in fluctuating environments

Under changing environments, the correlated fluctuation for an intrinsically coupled gene pair is a combination of intrinsic and extrinsic noise components. To establish a general framework for noise decomposition in this context, we refine Eq. (1) to explicitly separate the independent 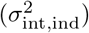 and coupled 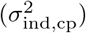 components of intrinsic fluctuations:

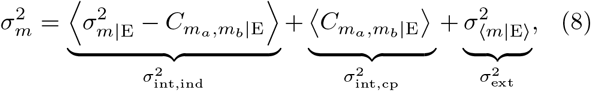

where 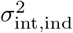 remains the uncorrelated component of the total fluctuation, while 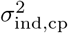 is defined as the average covariance between the nascent RNA levels of the two gene copies under fixed environmental conditions. Notably, both 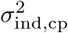 and 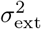 contribute to experimentally measured total covariance, i.e.:

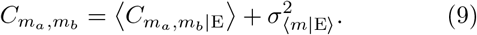

Hypothetically, 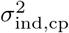 can be determined from Eq. (9) if the average expression level under every environmental condition (⟨*m*|E⟩) can be precisely measured.

Practically, however, determining the absolute values of ⟨*m*|E⟩ and 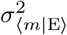 in fluctuating environments can be challenging. To distinguish 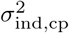 and 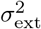 in a general sense, we notice that, if environmental fluctuations primarily affect gene-state transitions, then,

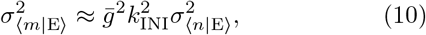

i.e., the extrinsic fluctuation is simply proportional to the variance of ON-state probabilities over fluctuating environments 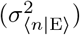, independent of their mean (⟨*n*⟩). In contrast, for *k*_ON_ & *k*_OFF_ ≲ 1, the coupled intrinsic fluctuation is a function of both ⟨*n*⟩ and 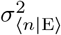 (Supplemental Material [27]):

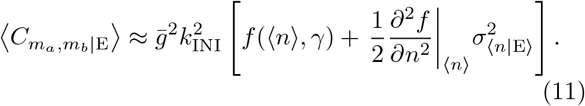

Therefore, the dependence of total covariance on ⟨*n*⟩ and 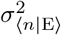 varies with the relative contributions of 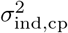 and 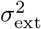 [Fig. 3(a)]. Specifically, for an independent gene pair 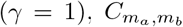 is solely a function of 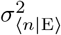. In contrast, for a fully coupled or anti-coupled gene pair (*γ* → ∞ or *γ* → 0), 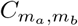 depends exclusively on ⟨*n*⟩ (Supplemental Material [27]). In all other cases, 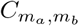 depends on both ⟨*n*⟩ and 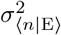, with its gradient offering insights into *γ*. In particular, with given ⟨*n*⟩, the partial slope of 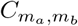 with respect to 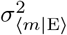 is simply

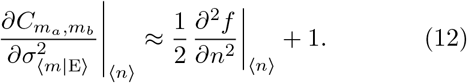

**FIG. 3.**
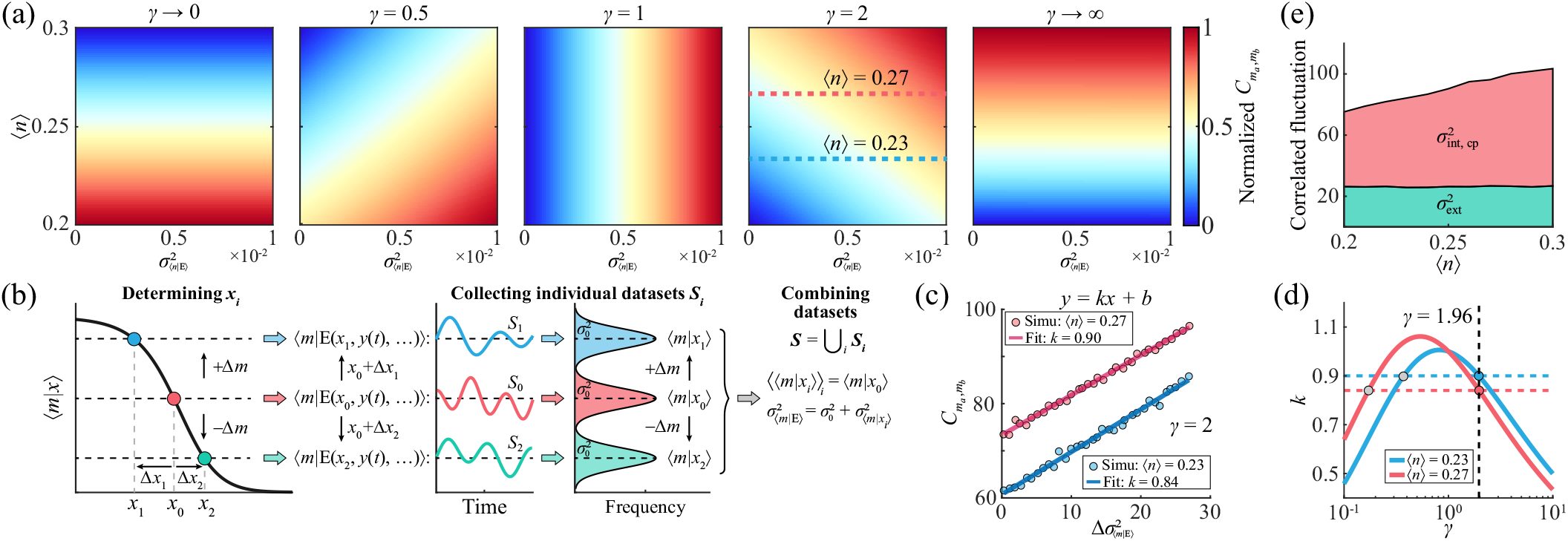
Distinguishing extrinsic and intrinsic components in correlated transcriptional fluctuations. (a) Total covariance of nascent RNA between gene copies as a function of ⟨*n*⟩ and 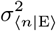 for different *γ* values. (b) Schematic of perturbing an external factor *x* above and below its original value to construct a combined dataset with increased 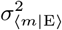 at fixed ⟨*m*⟩ (and ⟨*n*⟩). (c)Linear relationships between total covariance of nascent RNA and 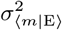 at different ⟨*n*⟩ values for the simulated data. (d) Estimation of *γ* from the slopes of the linear relationships in (c). (e) Noise decomposition of the simulated data in (c–d) as a function of ⟨*n*⟩ for fixed 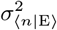. (b–e) The kinetic parameters are set as: *k*_ON_ = 0.01–0.2 min^−1^, *k*_OFF_ = 0.05 min^−1^, *k*_INI_ = 50 min^−1^, *T*_RES_ = 2 min, *g*(*τ*) = −*τ*, and *γ* = 2.

To apply this equation to experimental data, we note that, based on Eqs. (1), (5), and (7), ⟨*n* ⟩ can be estimated from the total noise of nascent RNA as

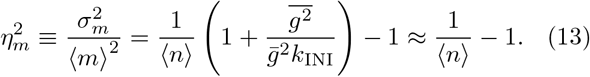

Here, 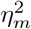, the squared coefficient of variation of nascent RNA, is experimentally measurable, and the approximation holds for *k*_INI_ ≫ 1, which is satisfied in most biological systems [13, 25, 51, 54–56]. Unlike ⟨*n*⟩, the absolute value of 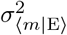 (or 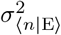), which may be influenced by multiple unknown or inaccessible external factors, is difficult to determine. However, if one of the *n* - regulating factors, denoted as *x*, is accessible, we may perturb it from the original value *x*_0_ to specific levels to collect datasets with distinct mean values (⟨*m*|*x*⟩) but comparable 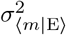 (Appendix B). By selecting a suitable set of perturbation levels {*x*_*i*_}, we can construct a combined dataset, whose average nascent RNA level ⟨⟨*m* |*x*_*i*_⟩ ⟩ _*i*_ matches that of the unperturbed dataset at *x*_0_ [Fig. 3(b)]. In contrast, 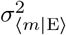 of the combined dataset increases with respect to the unperturbed one as

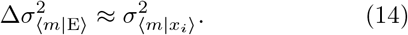

I.e., although the absolute value of 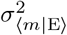 is unknown, we can quantitatively increase it while keeping ⟨*m*⟩ (and ⟨*n*⟩) fixed.

Building on these results, we propose a quantitative framework for estimating *γ*, 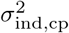, and 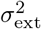 from experimental data through the following steps:

1. Determining ⟨*n*⟩: For any given experimental dataset, we can compute 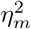 to estimate ⟨*n*⟩.
2. Measuring the partial slope of 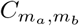: By carefully perturbing certain external factor, we can construct multiple datasets with varied 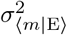at fixed ⟨*n*⟩. Computing the differences between these datasets in 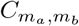 and 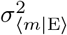 (using Eq. (14)) allows us to evaluate Eq. (12) at a given ⟨*n*⟩, which typically yields two possible solutions for *γ* [Fig. 3(c), see Supplemental Material [27]].
3. Determining *γ*: By repeating step 2 for different ⟨*n*⟩ values, we can identify the correct *γ* as the common solution across all cases [Fig. 3(d) and S2, see Supplemental Material [27]].
4. Decomposing 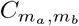: Once *γ* is determined, we can separate total covariance into 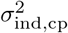 and 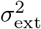 for any given ⟨*n*⟩ based on Eqs. (10)–(11) [Figs. 3(e), see Supplemental Material [27]].

Applying this framework to simulated data confirms its effectiveness and accuracy [Fig. 3(c–e)].

### Uncovering intrinsic couplings in Drosophila gene expression

To demonstrate the application of the above framework on experimental data, we examine the transcription of the *hunchback* (*hb*) gene in *Drosophila melanogaster* embryos [55] (Supplemental Material [27]). During early development, *hb* exhibits a logistic expression pattern along the anterior-posterior (AP) axis of the embryo, with its activation probability varying significantly across AP positions and *k*_ON_*T*_RES_ & *k*_OFF_*T*_RES_ ≲ 1 [24,51,57] [Fig. 4(a)]. In the later mitotic interphase of each cell cycle, there are two pairs of sister *hb* alleles in each nucleus. By quantifying nascent RNA at each *hb* allele in individual nuclei, we separately analyze correlated fluctuations for adjacent sister alleles and distant non-sister alleles, both previously assumed to be independent [24,51,57].

**FIG. 4.**
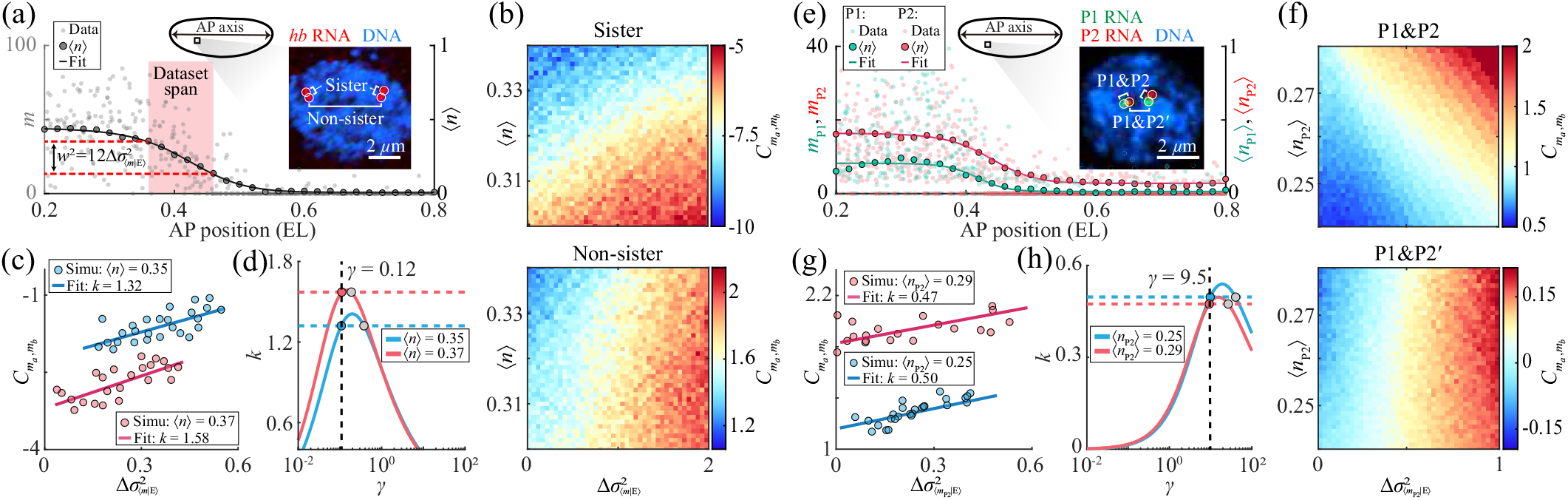
Decomposing correlated transcriptional fluctuations from experimental data. (a) Nascent RNA signal at individual *hb* genes in a single embryo as a function of AP position. Single-nucleus data are binned along the AP axis to estimate ⟨*n*⟩ for fitting to a logistic function. Each dataset with given ⟨*n*⟩ and 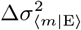 is constructed by uniformly sampling nuclei within a specific AP window (red region). Inset: confocal image of a nucleus showing nascent RNAs from two sets of sister *hb* alleles. (b) Total covariance of nascent RNA between sister alleles (upper) and between non-sister alleles (lower) as functions of ⟨*n*⟩ and 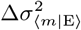. (c) Linear relationships between total covariance of nascent RNA and 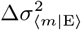 at different ⟨*n*⟩ values. (d) Estimation of *γ* between sister alleles from the slopes of the linear relationships in (c). (e) Nascent RNA signal at individual *hb* P1 and P2 (f) promoters in a single embryo as functions of AP position. Single-nucleus data are binned along the AP axis and to estimate ⟨*n*_P1_⟩ and ⟨*n*_P2_⟩ for fitting to logistic functions. Inset: confocal image of a nucleus showing nascent RNAs from P1 and P2. (f) Total covariance of nascent RNA between P1 and P2 on the same (upper) or different (lower) homologous alleles as functions of ⟨*n*_P2_⟩ and 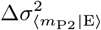. (g) Linear relationships between total covariance of nascent RNA and 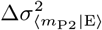 at different ⟨*n*_P2_⟩ values. (h) Estimation of *γ* between P1 and P2 from the slopes of the linear relationships in (g).

To apply our framework, we use the nuclear AP position *x* as a tuning factor for dataset construction. Specifically, by measuring the average *hb* expression level at each AP position (⟨*m* | *x*⟩), we uniformly sample single-nucleus data around each given position using varying *x*-ranges to construct datasets with distinct ⟨*n*⟩ and 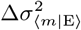 [Fig. 4(a), see Appendix C]. We find that the co-variance between non-sister alleles primarily varies with 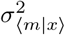, indicating independent transcription [Fig. 4(b)]. In contrast, the covariance between sister alleles depends on both ⟨*n*⟩ and 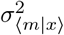, suggesting potential (anti-)coupling between them. Moreover, consistent with our theory, datasets with fixed ⟨*n*⟩ exhibit a linear relationship between 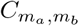 and 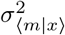 [Fig. 4(c)], whose slopes reveal a coupling strength of *γ* 0.12 and a negative 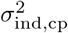 for sister alleles [Figs. 4(d) and S3(a), see Supplemental Material [27]]. This finding uncovers a previously unidentified anti-coupling, which may result from competition between adjacent sister alleles for transcription factor recruitment [14,15].

We then apply this strategy to investigate whether the two alternative promoters, P1 and P2, of the *hb* gene are coupled, a question unresolved in previous research due to the existence of extrinsic noise [55] [Fig. 4(e), Supplemental Material [27]]. These promoters are regulated by the same set of enhancers and therefore exhibit similar expression patterns (with different amplitudes) during early development [55]. By quantifying nascent RNA production from each promoter, we compare correlated fluctuations between P1 and P2 on either the same *hb* allele or different homologous alleles (denoted as P1 and P2’). We find that their 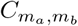 values exhibit distinct relationships with ⟨*n*⟩ and 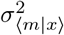 [Fig. 4(f)], suggesting potential coupling between P1 and P2 and independence between P1 and P2’. By extending our theory to describe non-identical gene pairs [Figs. S3(b–c), see Supplemental Material [27]], we estimate a coupling strength of *γ* ≈ 9.5 and a positive 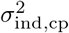 for P1 and P2 [Figs. 4(g–h) and S3(d)]. This previously unrecognized positive coupling is consistent with enhancer-shared promoter coupling observed in synthetic systems [20].

## Summary and outlook

In this Letter, we present a simple theoretical model to depict the stochastic nascent transcription of an intrinsically coupled gene pair under fluctuating environments. By analytically solving nascent RNA fluctuations, we develop a new noise decomposition framework to quantitatively distinguish correlated transcriptional fluctuations caused by intrinsic coupling from those induced by extrinsic variability, leveraging their distinct dependencies on the gene’s ON-state statistics. Applying it to single-cell transcription data from *Drosophila* embryos, we uncover couplings between sister alleles and between alternative promoters of an endogenous gene, both of which were challenging to identify in previous research.

Our findings broaden the classical picture of gene expression noise and provide a quantitative framework for deciphering intrinsic gene-gene interactions in fluctuating environments. While this study focuses on a specific *k*_ON_-coupling mechanism, our noise decomposition framework is broadly applicable to other coupling mechanisms (Supplemental Material [27]) and can be extended to investigate similar effects in other biochemical networks. Further generalizing this approach to analyze the temporal characteristics of correlated fluctuations among multiple genes will provide deeper insights into dynamic gene-gene interactions within complex gene regulatory networks.

## Acknowledgments

We thank Jingyao Wang for providing previously published imaging data, and Michael Levine and Bomyi Lim for their valuable discussions and feedback. This work was supported by the National Key R&D Program of China (grant no. 2021YFA0910702), the National Natural Science Foundation of China (grant no. 12474194, 11774225), and the Natural Science Foundation of Shanghai (grant no. 22ZR1434000). We gratefully acknowledge the imaging and computing resources provided by the Student Innovation Center at Shanghai Jiao Tong University, and sincerely thank Liuyin Fan for the dedicated management and support of these resources.

## Appendix A: Master equation analysis of the model

The stochastic dynamics of the system can be solved using either master equation or queueing theory approaches [23, 52, 53]. In this work,we adopt the master equation framework, as it offers a straightforward derivation of the statistical moments of nascent RNA. Specifically, by defining 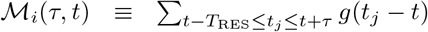 as the partial accumulation of nascent RNA signal over the history from *t* −*T*_RES_ to *t* + *τ*, we write the corresponding master equation as:

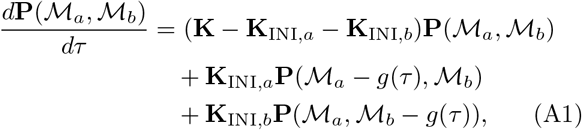

where 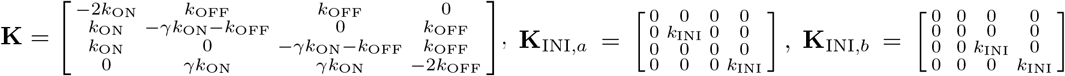, and 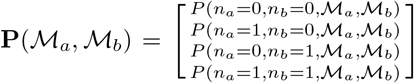. The probability distribution of the system state at time *t* corresponds to the solution of Eq. (A1) at *τ* = 0, i.e., **P**_*t*_(*m*_*a*_, *m*_*b*_) = **P**_*τ*=0_(ℳ_*a*_, ℳ_*b*_), given the initial marginal distribution of gene states [23]. Note that Eq. (A1) can be nondimensionalized by redefining *τ* as *τ/T*_RES_ and multiplying all kinetic rates by *T*_RES_. Without loss of generality, we set *T*_RES_ = 1 for this study.

Under a fixed environmental condition, the steady-state marginal distribution of gene state, 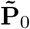, is easily solved from 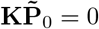, yielding Eq. (2). To further determine the fluctuations of *m*_*a*_ and *m*_*b*_, we apply characteristic function method to obtain Eq. (5) and (Supplemental Material [27]):

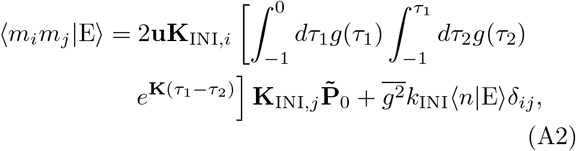

where 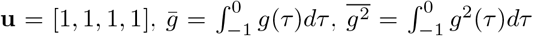 and *i, j* = *a* or *b*. Notably, **K** is negative semi-definite, whose eigenvalues are proportional to the magnitudes of *k*_ON_ and *k*_OFF_. Therefore, ⟨*m*_*i*_*m*_*j*_|E⟩ decays with the scale of gene-state transition rates.

Through a Taylor expansion of 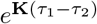 in Eq. (A2), we derive expressions for the nascent-RNA covariance and total variance as:

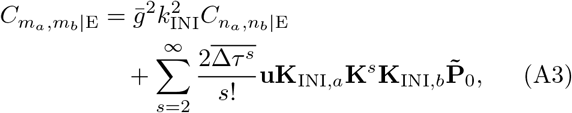

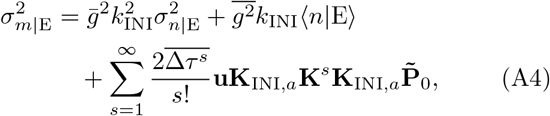

where 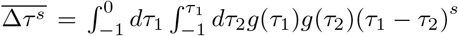. Since 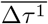 and 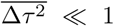 under slow gene-state transitions (Supplemental Material [27]), Eqs. (A3) and (A4) reduce to (6) and (7) without significant deviations until *k*_ON_ & *k*_OFF_ *>* 1 [Fig. 2(d) and S1(b)].

## Appendix B: Tuning 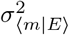 while keeping ⟨m⟩ fixed

Gene transcription in real biological systems is typically influenced by multiple external factors. While many of these factors remain unidentified or beyond experimental control, some may be sufficiently characterized to enable experimental modulation. Suppose we can manipulate a particular *k*_ON_- or *k*_OFF_-regulating factor, *x*, without influencing the fluctuations of other external factors (e.g., *y*(*t*)). By fixing *x* at a specific level, *x*_0_, we can collect a dataset *S*_0_ of *m*_*a*_ and *m*_*b*_ either from a time series of a single cell or from a snapshot of an ensemble of cells. The average gene transcription level in *S*_0_, denoted as ⟨*m* | *x*_0_⟩ ≡ ⟨⟨*m* | E(*x*_0_, *y*(*t*), …) ⟩⟩ _*t*_, is directly measurable. Due to the fluctuations of other external factors (and any trace fluctuation in *x*), the extrinsic variance of nascent RNA in *S*_0_, denoted as 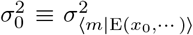, is nonzero [Fig. 3(b)].

Now consider applying a constant perturbation to *x* (i.e., *x*_1_ = *x*_0_ + Δ*x*_1_) to generate a new dataset *S*_1_ [Fig. 3(b)]. The average nascent RNA level will shift to ⟨*m* | *x*_1_⟩ accordingly. More importantly, the extrinsic variance of nascent RNA, to a first-order approximation, remains unchanged for constant Δ*x*_1_, i.e.,

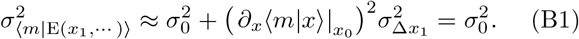

Therefore, if we collect multiple datavsets at a series of *x* values (*x*_*i*_, *i* = 1, 2, …) [Fig. 3(b)], the average nascent RNA level in the combined dataset *S* = ∪ _*i*_ *S*_*i*_ becomes:

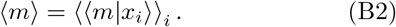

By knowing how ⟨*m*|*x*⟩ varies with *x*, we can select a suitable set of *x*_*i*_ around *x*_0_, so that *S* has the same ⟨*m*⟩ as *S*_0_. Meanwhile, the extrinsic variance of *m* in *S* satisfies:

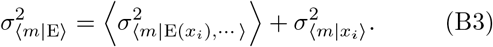

Since 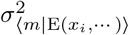 is roughly unchanged across individual datasets, the increase in 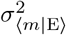 due to dataset combination can be approximated by 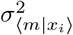 (Eq. 14). This strategy enables us to quantitatively tune 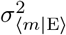 while keeping ⟨*m*⟩ (and ⟨*n*⟩) fixed.

## Appendix C: Constructing datasets for Drosophila embryos

During early *Drosophila* development, the *hb* gene is primarily expressed in the anterior half of the embryo, forming a logistic expression pattern along the AP axis. Previous studies have reported that such spatial variation in transcription is primarily driven by an AP-position (*x*) dependent regulation of *k*_ON_ [51,55]. Therefore, by grouping single-nucleus data across different *x* values, we can construct a series of datasets with systematically varied ⟨*n*⟩ and 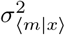 for covariance analysis. This proceeds as follows:

1. Measuring ⟨*n* | *x*⟩ as a function of nuclear position: By binning the single-nucleus data along the AP axis (bin size: 0.05 EL), we compute, for each bin, the average levels of nascent RNA (⟨*m* | *x*⟩) and ON-state probability (⟨*n* |*x*⟩, using Eq. (13)). Since these two quantities are typically proportional, we concentrate on ⟨*n* |*x*⟩ in the following description for simplicity. Here, we focus primarily on the AP-position range 0.20–0.60 embryo length (EL), where the per-allele nascent RNA level decreases systematically from high to low. Within this region, we smooth the trend of ⟨*n*|*x*⟩ with respect to *x* by fitting it to a logistic function 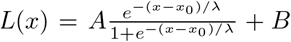, where *x*_0_, *λ, A*, and *B* are fitting parameters.
2. Selecting a set of *x* for dataset construction: To construct a combined dataset with desired values of ⟨*n*⟩ and 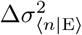, we first need to determine a set of *x* values for individual datasets. In principle, any *x*-set whose corresponding ⟨*n* | *x*⟩ values yield the target ⟨*n*⟩ and 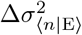 is acceptable. For simplicity, we consider a uniform distribution of ⟨*n* | *x*⟩ values centered at ⟨*n*⟩, with a width satisfying *w*^2^ = 12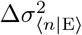. By discretizing this distribution (Δ*n* = 0.005), we obtain the corresponding *x* for each ⟨*n* | *x*⟩ via *x* = *L*^−1^ (⟨*n* | *x*⟩), yielding a set of *x* values.
3. Sampling single-nucleus data from each *x* to construct a combined dataset: For each *x* in the above set, we collect all single-nucleus data around it within a bin width of 0.005*/L*^′^(*x*) EL. This *x*-sensitive bin width is chosen to maintain a fixed range of ⟨*n* | *x*⟩ within each bin. Since the number of nuclei in each bin (denoted as *N*_*i*_) varies, we randomly sample an equal number of nuclei from each bin (*N* = min(*N*_*i*_)) to construct a combined dataset. This guarantees balanced contributions from each AP position in the final dataset.

