## Supplemental Material for "Extrinsic Noise or Intrinsic Coupling: Dissecting Correlated Fluctuations in Gene Transcription"

### Table of Contents:

**1. Model assumptions ... 3**

**2. Numerical methods for solving the model ... 5**

**3. Deriving the steady-state properties of gene-state distribution ... 6**

**4. Deriving the moments of nascent RNA distribution ... 7**

4.1 General results ... 7

4.2 Slow gene-state transitions ( $k_{\mathrm{ON}}T_{\mathrm{RES}}, k_{\mathrm{OFF}}T_{\mathrm{RES}} \ll 1$ or $k_{\mathrm{ON}}T_{\mathrm{RES}}, k_{\mathrm{OFF}}T_{\mathrm{RES}} \lesssim 1$) ... 9

4.3 Fast gene-state transitions ( $k_{\mathrm{ON}}T_{\mathrm{RES}}, k_{\mathrm{OFF}}T_{\mathrm{RES}} \gg 1$ ) ... 11

**5. Deriving the extrinsic fluctuation of nascent RNA ... 12**

**6. Deriving the coupled intrinsic fluctuation in noisy environments ... 13**

6.1 Zeroth-order results ... 13

6.2 Correction for $k_{\mathrm{ON}}T_{\mathrm{RES}}, k_{\mathrm{OFF}}T_{\mathrm{RES}} \lesssim 1$ ... 14

**7. Deriving the total correlated fluctuation for $\gamma = 0, 1,$ and $\infty$ ... 15**

**8. Deriving the total transcriptional fluctuation for a single gene ... 16**

8.1 Zeroth-order results ... 16

8.2 Correction for $k_{\mathrm{ON}}T_{\mathrm{RES}}, k_{\mathrm{OFF}}T_{\mathrm{RES}} \lesssim 1$ ... 16

**9. Alternative methods for determining $\gamma$ ... 17**

**10. Separating extrinsic fluctuation and coupled intrinsic fluctuation ... 18**

**11. Experimental data extraction ... 18**

**12. Model extension for non-identical gene copies ... 20**

**13. Generalization of the framework to diverse coupling mechanisms ... 23**

13.1. TF-sharing mechanism ... 23

13.2. $k_{\mathrm{ON}}$-coupling in prokaryotes with co-transcriptional degradation ... 24

13.3. $k_{\mathrm{OFF}}$-coupling mechanism ... 25

13.4. Coupled multi-state models ... 26

13.4.1. Coupled "OFF-OFF-ON" model ... 27

13.4.2. Coupled "OFF-ON-ON" model ... 29

13.5. RNAP-sharing mechanism ... 30

13.6. Feedback mechanisms ... 32

13.6.1. Autoregulation ... 33

13.6.2. Toggle switch ... 34

#### Supplemental Material

**Supplemental Material**

#### 1. Model assumptions

In this study, we focus on modeling nascent RNA, whose lifetime typically spans only a few minutes. This short lifetime acts as a natural filter that suppresses the influence of slow coupling processes, such as protein-mediated feedback regulations[1,2], thereby emphasizing on more direct coupling mechanisms. In practice, nascent RNA signals can be captured using short-lived reporter systems such as the MS2 system in live-cell imaging[3–5] or smFISH in fixed-cell imaging[6–8]. Alternatively, short-lived pre-RNA signals detected via single-cell RNA sequencing can be used[9].

We model two sister gene copies within a single eukaryotic cell based on the standard two-state telegraph model for nascent transcription[10,11]. This model describes transcription process of each gene copy in four steps: gene activation/inactivation, transcription initiation in the active (ON) state, nascent RNA synthesis (elongation), and release. Gene state transitions and transcription initiation are modeled as Poisson processes, with rates $k_{\mathrm{ON}}$, $k_{\mathrm{OFF}}$, and $k_{\mathrm{INI}}$, respectively[10–14]. In contrast, nascent RNA elongation and release are treated as deterministic processes, with a constant speed $V_{\mathrm{EL}}$ and a post-elongation residence duration $T_{\mathrm{S}}$[15,16], leading to a single nascent RNA lifetime of $T_{\mathrm{RES}} = L / V_{\mathrm{EL}} + T_{\mathrm{S}}$. These assumptions are widely used in previous studies of bursty single-cell transcription kinetics[11–13,17,18], and can be readily extended to incorporate more complex kinetics of gene activation/inactivation, transcription initiation, and RNA release[7,11]. For example, while we assume two identical gene copies with the same kinetic parameters here, the model can be easily adapted to describe non-identical gene copies with distinct kinetic parameters (see **Section 12**).

To model the coupling between gene copies, particularly at the level of gene activation, we assume that if one gene copy switches to the ON state, the activation rate of its sister copy increases by a factor $\gamma$, i.e., $k_{\mathrm{ON}} \to \gamma k_{\mathrm{ON}}$. This simple assumption fits the mechanisms of enhancer-shared gene activation[19–21], and serves as a prototype for modeling coupling mechanisms at other gene expression steps, such as gene inactivation, transcription initiation, and translation. Notably, $\gamma > 1$ and $0 < \gamma < 1$ correspond to positive and negative couplings, respectively, highlighting the versatility of our model.

To accurately model the experimentally observed nascent RNA signals, we assume that each nascent RNA contributes a non-integer signal based on its length (or its initiation time relative to the observation moment). Following previous studies[6,11,18], we define a contribution function $g(\tau)$ to describe this quantitative relationship, with $\tau$ denoting the relative initiation time ($-T_{\mathrm{RES}} \le \tau \le 0$). Note that $g(\tau)$ is a general function, whose specific form depends on the RNA labeling method. For example, if only the 5' end of the RNA is labeled, then $g(\tau) = 1$. In contrast, if the entire RNA is uniformly labeled, then $g(\tau) = -\tau / T_{\mathrm{RES}}$. The contribution functions used in this study are summarized in **Table S1**.

The influence of fluctuating environments on gene transcription can be modeled by allowing all kinetic parameters to vary over time and across cells. Here, based on previous research[22,23], we assume that environmental fluctuations occur on a time scale slower than that of nascent transcription ($T_{\mathrm{RES}}$). Under this premise, these fluctuations can be represented by a static probability distribution $p(\mathrm{E})$ of kinetic parameters $\mathrm{E} = [k_{\mathrm{ON}}, k_{\mathrm{OFF}}, k_{\mathrm{INI}}, T_{\mathrm{RES}}, \gamma]$, while the system state is approximated as a mixture of steady states for all possible E[22]. As an example, this adiabatic assumption applies to a special case where the kinetic parameters are cell-cycle dependent. Given that the lifetime of nascent RNA is typically much shorter than the duration of a cell cycle, the kinetic rates can be considered quasi-static over the duration of nascent RNA production. Meanwhile, the cell-cycle-induced fluctuations can be treated as a static probability distribution of kinetic rates, i.e., they function as a regular form of extrinsic noise. Furthermore, based on prior studies, we assume that $k_{\mathrm{ON}}$ and $k_{\mathrm{OFF}}$ are the primary targets of environmental fluctuations[23,24].

For rapid environmental fluctuations, however, the system may never reach (near-)steady state. In this case, our approach may be extended by replacing the conditioning on a static E with conditioning on the entire history of E[25,26].

Note that our model primarily focuses on eukaryotic gene expression, where gene copy number remains relatively stable (two or four copies per cell) in typical cases. In contrast, in prokaryotes such as *E. coli*, gene copy number can vary considerably due to the presence of multiple active replication forks during rapid growth[27]. This variability introduces additional uncertainty into transcriptional dynamics, complicating the modeling of gene copy interactions. Furthermore, prokaryotic transcription often involves co-transcriptional degradation of nascent RNA, requiring a modified modeling framework to accurately capture the RNA dynamics[28,29]. Therefore, even without considering gene copy number variation, extending our theoretical framework to bacterial gene expression requires incorporating this sophisticated degradation mechanism into the model. In **Section 13.2** of this document, we show through theoretical and computational analyses that our noise decomposition framework remains valid for this modified model.

#### 2. Numerical methods for solving the model

Stochastic simulation of the model was performed in MATLAB 2021a following the Gillespie algorithm[6,11,30]. The kinetic parameters for simulations are presented in the corresponding figure legends or **Supplemental Material** sections. In **Figs. 3(c, e)**, to simulate gene transcription under fluctuation environments with specified $\langle n \rangle$ and $\sigma^2_{\langle n|\mathrm{E}\rangle}$, we first construct a discretized uniform probability distribution for $\langle n \mid \mathrm{E} \rangle$ ($\Delta n = 0.0005$), centered at $\langle n \rangle$ with a variance of $\sigma^2_{\langle n|\mathrm{E}\rangle}$. For each $\langle n \mid \mathrm{E} \rangle$ bin in this distribution, we determine the corresponding $k_{\mathrm{ON}}$ values to perform 100,000 simulations. Pooling the results from all $\langle n \mid \mathrm{E} \rangle$ bins yields a dataset for the specified $\langle n \rangle$ and $\sigma^2_{\langle n|\mathrm{E}\rangle}$.

**Supplemental Material**

**3. Deriving the steady-state properties of gene-state distribution**

The steady-state marginal distribution of gene states under a given environmental condition is solved from the equation $\mathbf{K}\tilde{\mathbf{P}}_0 = 0$ as:

$$\tilde{\mathbf{P}}_0 \equiv \begin{bmatrix} P(n_a = 0, n_b = 0 \mid \mathrm{E}) \\ P(n_a = 1, n_b = 0 \mid \mathrm{E}) \\ P(n_a = 0, n_b = 1 \mid \mathrm{E}) \\ P(n_a = 1, n_b = 1 \mid \mathrm{E}) \end{bmatrix} = \frac{1}{1 + 2\left(k_{\mathrm{ON}} / k_{\mathrm{OFF}}\right) + \gamma\left(k_{\mathrm{ON}} / k_{\mathrm{OFF}}\right)^2} \begin{bmatrix} 1 \\ k_{\mathrm{ON}} / k_{\mathrm{OFF}} \\ k_{\mathrm{ON}} / k_{\mathrm{OFF}} \\ \gamma\left(k_{\mathrm{ON}} / k_{\mathrm{OFF}}\right)^2 \end{bmatrix}. \tag{S1}$$

From this distribution, we can easily write down the covariance (**Eq. (5)**), the correlation coefficient:

$$\rho_{n_a, n_b \mid \mathrm{E}} = \frac{\gamma - 1}{\left(\gamma + \dfrac{k_{\mathrm{OFF}}}{k_{\mathrm{ON}}}\right)\left(1 + \dfrac{k_{\mathrm{ON}}}{k_{\mathrm{OFF}}}\right)}, \tag{S2}$$

and the ON-state probability of a gene copy:

$$\langle n_a \mid \mathrm{E} \rangle \equiv P(n_a = 1 \mid \mathrm{E}) = P(n_b = 1 \mid \mathrm{E}) = \frac{\left(k_{\mathrm{ON}} / k_{\mathrm{OFF}}\right) + \gamma\left(k_{\mathrm{ON}} / k_{\mathrm{OFF}}\right)^2}{1 + 2\left(k_{\mathrm{ON}} / k_{\mathrm{OFF}}\right) + \gamma\left(k_{\mathrm{ON}} / k_{\mathrm{OFF}}\right)^2}, \tag{S3}$$

all of which depend on $\dfrac{k_{\mathrm{ON}}}{k_{\mathrm{OFF}}}$ and $\gamma$. Therefore, we can rewrite the covariance (and the correlation coefficient) as a function of $\langle n_a \mid \mathrm{E} \rangle$ (abbreviated as $n_{\mathrm{E}}$) and $\gamma$:

$$\mathrm{Cov}(n_a, n_b \mid \mathrm{E}) = f(n_{\mathrm{E}}, \gamma) = (\gamma - 1)\left[\frac{(1 - n_{\mathrm{E}})\left(2n_{\mathrm{E}} - 1 + \sqrt{4(\gamma - 1)n_{\mathrm{E}}(1 - n_{\mathrm{E}}) + 1}\right)}{2\gamma(1 - n_{\mathrm{E}}) + 2n_{\mathrm{E}} - 1 + \sqrt{4(\gamma - 1)n_{\mathrm{E}}(1 - n_{\mathrm{E}}) + 1}}\right]^2, \tag{S4}$$

and $\rho_{n_a, n_b \mid \mathrm{E}} = \dfrac{f(n_{\mathrm{E}}, \gamma)}{n_{\mathrm{E}}(1 - n_{\mathrm{E}})}$. For a given $\gamma$, both the correlation coefficient and the covariance show extreme values $\rho_{n_a, n_b \mid \mathrm{E}} = \dfrac{\sqrt{\gamma} - 1}{\sqrt{\gamma} + 1}$ and $\mathrm{Cov}(n_a, n_b \mid \mathrm{E}) = \dfrac{1}{4}\dfrac{\sqrt{\gamma} - 1}{\sqrt{\gamma} + 1}$ at $k_{\mathrm{ON}} / k_{\mathrm{OFF}} = \gamma^{-1/2}$, which corresponds to $\langle n \mid \mathrm{E} \rangle = 0.5$.

#### Supplemental Material

##### 4. Deriving the moments of nascent RNA distribution

###### 4.1 General results

By defining the characteristic function $\mathbf{\Psi}(\omega_a,\omega_b) \equiv \int_0^\infty d\mathcal{m}_a \int_0^\infty d\mathcal{m}_b e^{i\mathcal{m}_a\omega_a} e^{i\mathcal{m}_b\omega_b} \mathbf{P}(\mathcal{m}_a,\mathcal{m}_b)$, we can convert the master equation (**Eq. (3)**) to:

$$\frac{d\mathbf{\Psi}(\omega_a,\omega_b)}{d\tau} = \left[\mathbf{K} + (e^{i\omega_a g(\tau)} - 1)\mathbf{K}_{\mathrm{INI},a} + (e^{i\omega_b g(\tau)} - 1)\mathbf{K}_{\mathrm{INI},b}\right]\mathbf{\Psi}(\omega_a,\omega_b). \tag{S5}$$

Since the moments of $m_a$ and $m_b$ under a given environmental condition satisfy

$$\left\langle m_a^M m_b^N \mid \mathrm{E}\right\rangle = \mathbf{u}\cdot(-i)^{M+N} \left.\frac{\partial^{M+N}\mathbf{\Psi}_{\tau=0}(\omega_a,\omega_b)}{\partial\omega_a^M \partial\omega_b^N}\right|_{\omega_a=0,\omega_b=0}, \tag{S6}$$

with $\mathbf{u}=[1,1,1,1]$, they can be derived by taking derivatives of **Eq. (S6)** with respect to $\omega_a$ and $\omega_b$, i.e.:

$$\frac{d}{d\tau}\left(\left.\frac{\partial^{M+N}\mathbf{\Psi}(\omega_a,\omega_b)}{\partial\omega_a^M\partial\omega_b^N}\right|_{\substack{\omega_a=0,\\ \omega_b=0}}\right) = \mathbf{K}\left.\frac{\partial^{M+N}\mathbf{\Psi}(\omega_a,\omega_b)}{\partial\omega_a^M\partial\omega_b^N}\right|_{\substack{\omega_a=0,\\ \omega_b=0}} + \mathbf{K}_{\mathrm{INI},a}\sum_{j=0}^{M-1}\binom{M}{j}(ig)^{M-j}\left.\frac{\partial^{j+N}\mathbf{\Psi}(\omega_a,\omega_b)}{\partial\omega_a^j\partial\omega_b^N}\right|_{\substack{\omega_a=0,\\ \omega_b=0}}$$
$$+\mathbf{K}_{\mathrm{INI},b}\sum_{k=0}^{N-1}\binom{N}{k}(ig)^{N-k}\left.\frac{\partial^{M+k}\mathbf{\Psi}(\omega_a,\omega_b)}{\partial\omega_a^M\partial\omega_b^k}\right|_{\substack{\omega_a=0,\\ \omega_b=0}}. \tag{S7}$$

This is a set of first-order linear differential equations with respect to $\tau$, whose solutions at $\tau=0$ satisfy:

$$\left\langle m_a^M m_b^N \mid \mathrm{E}\right\rangle = \mathbf{u}\cdot\sum_{I,J=1}^{M,N}\underbrace{\sum_{0=r_0<r_i<r_I=M}}_{i=1,\cdots,I-1}\underbrace{\sum_{0=s_0<s_j<s_J=N}}_{j=1,\cdots,J-1}\sum_{\sigma\in\mathrm{Interleave}(I,J)}\int_{-T_{\mathrm{RES}}}^{0}d\tau_1\cdots$$
$$\times\int_{-T_{\mathrm{RES}}}^{\tau_{I+J-1}}d\tau_{I+J}\mathcal{T}\left[\prod_{k=1}^{I+J}\binom{C_{\sigma(k)}}{C'_{\sigma(k)-1}}g(\tau_k)^{C_{\sigma(k)}-C'_{\sigma(k)-1}}\mathbf{W}_{H(\sigma(k)-I-1)}(\tau_k)\right]e^{-\mathbf{K}\tau_{I+J}}\mathbf{\Psi}_0, \tag{S8}$$

where $\mathrm{Interleave}(I,J)$ represents a set consisting of all permutations of $\{1,\cdots I+J\}$ that preserve the relative order from 1 to *I* and from *I*+1 to *I*+*J*, with individual elements denoted by $\sigma$, $\mathcal{T}$ is the time-ordering operator, $C=\{r_1,\cdots,r_M,s_1,\cdots,s_N\}$, $\sigma(k)$ is the *k*'s element of a specific $\sigma$, $C_{\sigma(k)}$ is the $\sigma(k)$'s element of $C$, $C'_{\sigma(k)-1}=C_{\sigma(k)-1}(1-\delta_{\sigma(k),1})(1-\delta_{\sigma(k),I+1})$, $H(x)$ is the Heaviside step function, $\mathbf{W}_0(\tau)=\mathbf{W}_a(\tau)=e^{-\mathbf{K}\tau}\mathbf{K}_{\mathrm{INI},a}e^{\mathbf{K}\tau}$, $\mathbf{W}_1(\tau)=\mathbf{W}_b(\tau)=e^{-\mathbf{K}\tau}\mathbf{K}_{\mathrm{INI},b}e^{\mathbf{K}\tau}$, and $\mathbf{\Psi}_0=\mathbf{\Psi}_{\tau=-T_{\mathrm{RES}}}(0,0)=\tilde{\mathbf{P}}_0$.

**Supplemental Material**

Specifically, the first-order moment of $m_a$ (and $m_b$) is derived to be

$$\begin{aligned}\langle m_a \mid \mathrm{E}\rangle &= \mathbf{u}\left[\int_{-T_{\mathrm{RES}}}^{0} d\tau g(\tau)\mathbf{W}_a(\tau)e^{-\mathbf{K}\tau}\right]\mathbf{\Psi}_0 = \left[\int_{-T_{\mathrm{RES}}}^{0} d\tau g(\tau)\mathbf{u}e^{-\mathbf{K}\tau}\mathbf{K}_{\mathrm{INI},a}\right]\mathbf{\Psi}_0 \\ &= \left[\int_{-T_{\mathrm{RES}}}^{0} d\tau g(\tau)\right]\mathbf{u}\mathbf{K}_{\mathrm{INI},a}\mathbf{\Psi}_0 = k_{\mathrm{INI}}T_{\mathrm{RES}}\overline{g}\langle n_a \mid \mathrm{E}\rangle,\end{aligned} \tag{S9}$$

where $\overline{g} = \frac{1}{T_{\mathrm{RES}}}\int_{-T_{\mathrm{RES}}}^{0} g(\tau)d\tau$. Here, the simplification $\mathbf{u}e^{-\mathbf{K}\tau} = \mathbf{u}$ arises from the fact that each column of $\mathbf{K}$ sums to zero. Similarly, the second-order moments of $m_a$ and $m_b$ are derived to be

$$\begin{aligned}\langle m_a^2 \mid \mathrm{E}\rangle &= \mathbf{u}\left[2\int_{-T_{\mathrm{RES}}}^{0} d\tau_1 g(\tau_1)\mathbf{W}_a(\tau_1)\int_{-T_{\mathrm{RES}}}^{\tau_1} d\tau_2 g(\tau_2)\mathbf{W}_a(\tau_2)e^{-\mathbf{K}\tau_2} + \int_{-T_{\mathrm{RES}}}^{0} d\tau g^2(\tau)\mathbf{W}_a(\tau)e^{-\mathbf{K}\tau}\right]\mathbf{\Psi}_0 \\ &= \left[2\int_{-T_{\mathrm{RES}}}^{0} d\tau_1 g(\tau_1)\int_{-T_{\mathrm{RES}}}^{\tau_1} d\tau_2 g(\tau_2)\mathbf{u}e^{-\mathbf{K}\tau_1}\mathbf{K}_{\mathrm{INI},a}e^{\mathbf{K}(\tau_1-\tau_2)}\mathbf{K}_{\mathrm{INI},a} + \int_{-T_{\mathrm{RES}}}^{0} d\tau g^2(\tau)\mathbf{u}e^{-\mathbf{K}\tau}\mathbf{K}_{\mathrm{INI},a}\right]\mathbf{\Psi}_0 \\ &= 2\mathbf{u}\mathbf{K}_{\mathrm{INI},a}\mathbf{Q}\left[\int_{-T_{\mathrm{RES}}}^{0} d\tau_1 g(\tau_1)\int_{-T_{\mathrm{RES}}}^{\tau_1} d\tau_2 g(\tau_2)e^{\mathbf{K}_0(\tau_1-\tau_2)}\right]\mathbf{Q}^{-1}\mathbf{K}_{\mathrm{INI},a}\mathbf{\Psi}_0 + k_{\mathrm{INI}}T_{\mathrm{RES}}\overline{g^2}\langle n_a \mid \mathrm{E}\rangle,\end{aligned} \tag{S10}$$

$$\begin{aligned}\langle m_a m_b \mid \mathrm{E}\rangle &= \mathbf{u}\left\{\int_{-T_{\mathrm{RES}}}^{0} g(\tau_1)\int_{-T_{\mathrm{RES}}}^{\tau_1} g(\tau_2)\left[\mathbf{W}_a(\tau_1)\mathbf{W}_b(\tau_2) + \mathbf{W}_b(\tau_1)\mathbf{W}_a(\tau_2)\right]e^{-\mathbf{K}\tau_2}d\tau_1 d\tau_2\right\}\mathbf{\Psi}_0 \\ &= \left\{\int_{-T_{\mathrm{RES}}}^{0} d\tau_1 g(\tau_1)\int_{-T_{\mathrm{RES}}}^{\tau_1} d\tau_2 g(\tau_2)\mathbf{u}e^{-\mathbf{K}\tau_1}\left[\mathbf{K}_{\mathrm{INI},a}e^{\mathbf{K}(\tau_1-\tau_2)}\mathbf{K}_{\mathrm{INI},b} + \mathbf{K}_{\mathrm{INI},b}e^{\mathbf{K}(\tau_1-\tau_2)}\mathbf{K}_{\mathrm{INI},a}\right]\right\}\mathbf{\Psi}_0 \\ &= 2\int_{-T_{\mathrm{RES}}}^{0} d\tau_1 g(\tau_1)\int_{-T_{\mathrm{RES}}}^{\tau_1} d\tau_2 g(\tau_2)\mathbf{u}\mathbf{K}_{\mathrm{INI},a}e^{\mathbf{K}(\tau_1-\tau_2)}\mathbf{K}_{\mathrm{INI},b}\mathbf{\Psi}_0 \\ &= 2\mathbf{u}\mathbf{K}_{\mathrm{INI},a}\mathbf{Q}\left[\int_{-T_{\mathrm{RES}}}^{0} d\tau_1 g(\tau_1)\int_{-T_{\mathrm{RES}}}^{\tau_1} d\tau_2 g(\tau_2)e^{\mathbf{K}_0(\tau_1-\tau_2)}\right]\mathbf{Q}^{-1}\mathbf{K}_{\mathrm{INI},b}\mathbf{\Psi}_0,\end{aligned} \tag{S11}$$

where $\overline{g^2} = \frac{1}{T_{\mathrm{RES}}}\int_{-T_{\mathrm{RES}}}^{0} g^2(\tau)d\tau$. The first term on the right-hand side of **Eq. (S10)** denotes the variance caused by transitions between gene states, while the second term accounts for the contribution from Poissonian transcription initiation during the ON state[11]. In contrast, the covariance between the two gene copies' expression levels is entirely driven by the covariance between their gene states.

In the above derivation, we have used diagonalization to simplify the computation of the integral, with $\mathbf{Q}$ and $\mathbf{K}_0 = \mathbf{diag}(k_i)$ denoting eigenvector and eigenvalue matrices of $\mathbf{K}$, respectively, i.e.,

$$k_i = 0, -k_{\mathrm{OFF}} - \gamma k_{\mathrm{ON}}, -\frac{3k_{\mathrm{OFF}} + (2+\gamma)k_{\mathrm{ON}} \pm \sqrt{k_{\mathrm{OFF}}^2 + (6\gamma-4)k_{\mathrm{ON}}k_{\mathrm{OFF}} + (\gamma-2)^2 k_{\mathrm{ON}}^2}}{2}, \tag{S12}$$

$$\mathbf{Q}=\begin{bmatrix} \dfrac{k_{\mathrm{OFF}}^2}{\gamma k_{\mathrm{ON}}^2} & 0 & \dfrac{-k_3-2k_{\mathrm{OFF}}-\gamma k_{\mathrm{ON}}}{\gamma k_{\mathrm{ON}}} & \dfrac{-k_4-2k_{\mathrm{OFF}}-\gamma k_{\mathrm{ON}}}{\gamma k_{\mathrm{ON}}} \\ \dfrac{k_{\mathrm{OFF}}}{\gamma k_{\mathrm{ON}}} & -1 & \dfrac{k_3+2k_{\mathrm{OFF}}}{2\gamma k_{\mathrm{ON}}} & \dfrac{k_4+2k_{\mathrm{OFF}}}{2\gamma k_{\mathrm{ON}}} \\ \dfrac{k_{\mathrm{OFF}}}{\gamma k_{\mathrm{ON}}} & 1 & \dfrac{k_3+2k_{\mathrm{OFF}}}{2\gamma k_{\mathrm{ON}}} & \dfrac{k_4+2k_{\mathrm{OFF}}}{2\gamma k_{\mathrm{ON}}} \\ 1 & 0 & 1 & 1 \end{bmatrix}. \tag{S13}$$

We can see that one of the eigenvalues is zero, while the others are all negative and proportional to the magnitude of $k_{\mathrm{ON}}$ and $k_{\mathrm{OFF}}$ (It can be expressed as $k_{\mathrm{OFF}}$ multiplied by a function of $\dfrac{k_{\mathrm{ON}}}{k_{\mathrm{OFF}}}$ and $\gamma$). In contrast, $\mathbf{Q}$ depends only on $\dfrac{k_{\mathrm{ON}}}{k_{\mathrm{OFF}}}$ and $\gamma$. These results indicate that, for given values of $\dfrac{k_{\mathrm{ON}}}{k_{\mathrm{OFF}}}$ and $\gamma$, $\langle m_a m_b \mid \mathrm{E}\rangle$ and the first term in $\langle m_a^2 \mid \mathrm{E}\rangle$ both decay with gene-state transition rates.

While the above formulas allow for the derivation of analytical expressions for specific contribution functions, such as $g=1$ or $g=-\tau/T_{\mathrm{RES}}$, these expressions are typically too lengthy to present here. In the following, we analyze two specific cases.

##### 4.2 Slow gene-state transitions ($k_{\mathrm{ON}}T_{\mathrm{RES}}, k_{\mathrm{OFF}}T_{\mathrm{RES}} \ll 1$ or $k_{\mathrm{ON}}T_{\mathrm{RES}}, k_{\mathrm{OFF}}T_{\mathrm{RES}} \lesssim 1$)

For small $k_{\mathrm{ON}}T_{\mathrm{RES}}$ and $k_{\mathrm{OFF}}T_{\mathrm{RES}}$, **Eqs. (S10)-(S11)** can be rewritten as a series expansion of $e^{\mathbf{K}(\tau_1-\tau_2)}$:

$$\begin{aligned}\langle m_a^2 \mid \mathrm{E}\rangle &= \left\{\sum_{s=0}^{\infty}\frac{2\mathbf{u}\mathbf{K}_{\mathrm{INI},a}\mathbf{K}^s\mathbf{K}_{\mathrm{INI},a}\mathbf{\Psi}_0}{s!}\left[\int_{-T_{\mathrm{RES}}}^{0}d\tau_1\int_{-T_{\mathrm{RES}}}^{\tau_1}d\tau_2 g(\tau_1)g(\tau_2)(\tau_1-\tau_2)^s\right]\right\}+k_{\mathrm{INI}}T_{\mathrm{RES}}\overline{g^2}\langle n_a \mid \mathrm{E}\rangle \\ &= \mathbf{u}\mathbf{K}_{\mathrm{INI},a}^2\mathbf{\Psi}_0 T_{\mathrm{RES}}^2\bar{g}^2 + 2\mathbf{u}\mathbf{K}_{\mathrm{INI},a}\mathbf{K}\mathbf{K}_{\mathrm{INI},a}\mathbf{\Psi}_0\left[\int_{-T_{\mathrm{RES}}}^{0}d\tau_1\int_{-T_{\mathrm{RES}}}^{\tau_1}d\tau_2 g(\tau_1)g(\tau_2)(\tau_1-\tau_2)\right] \\ &\quad + \mathbf{u}\mathbf{K}_{\mathrm{INI},a}\mathbf{K}^2\mathbf{K}_{\mathrm{INI},a}\mathbf{\Psi}_0\left[\int_{-T_{\mathrm{RES}}}^{0}d\tau_1\int_{-T_{\mathrm{RES}}}^{\tau_1}d\tau_2 g(\tau_1)g(\tau_2)(\tau_1-\tau_2)^2\right]+\cdots+k_{\mathrm{INI}}T_{\mathrm{RES}}\overline{g^2}\langle n_a \mid \mathrm{E}\rangle,\end{aligned} \tag{S14}$$

$$
\begin{aligned}
\left\langle m_a m_b \mid \mathrm{E}\right\rangle &= \sum_{s=0}^{\infty} \frac{2\mathbf{uK}_{\mathrm{INI},a}\mathbf{K}^s\mathbf{K}_{\mathrm{INI},b}\boldsymbol{\Psi}_0}{s!}\left[\int_{-T_{\mathrm{RES}}}^{0} d\tau_1 \int_{-T_{\mathrm{RES}}}^{\tau_1} d\tau_2 g(\tau_1)g(\tau_2)(\tau_1-\tau_2)^s\right] \\
&= \mathbf{uK}_{\mathrm{INI},a}\mathbf{K}_{\mathrm{INI},b}\boldsymbol{\Psi}_0 T_{\mathrm{RES}}^2 \bar{g}^2 + \cancel{2\mathbf{uK}_{\mathrm{INI},a}\mathbf{K}\mathbf{K}_{\mathrm{INI},b}\boldsymbol{\Psi}_0\left[\int_{-T_{\mathrm{RES}}}^{0} d\tau_1 \int_{-T_{\mathrm{RES}}}^{\tau_1} d\tau_2 g(\tau_1)g(\tau_2)(\tau_1-\tau_2)\right]} \\
&\quad + \mathbf{uK}_{\mathrm{INI},a}\mathbf{K}^2\mathbf{K}_{\mathrm{INI},b}\boldsymbol{\Psi}_0\left[\int_{-T_{\mathrm{RES}}}^{0} d\tau_1 \int_{-T_{\mathrm{RES}}}^{\tau_1} d\tau_2 g(\tau_1)g(\tau_2)(\tau_1-\tau_2)^2\right] + \cdots,
\end{aligned}
\tag{S15}
$$

where the $s$th-order term is proportional to $k_{\mathrm{OFF}}^s$. Notably, the first-order term in $\left\langle m_a m_b \mid \mathrm{E}\right\rangle$ cancels out because $\mathbf{uK}_{\mathrm{INI},a}\mathbf{K}\mathbf{K}_{\mathrm{INI},b}\boldsymbol{\Psi}_0 = 0$.

For extremely slow gene-state transitions with $k_{\mathrm{ON}}T_{\mathrm{RES}},\ k_{\mathrm{OFF}}T_{\mathrm{RES}} \ll 1$, the second-order moments are entirely determined by the zeroth-order terms in **Eqs. (S14)-(S15)**:

$$
\left\langle m_a^2 \mid \mathrm{E}\right\rangle = \mathbf{uK}_{\mathrm{INI},a}^2\left[\int_{-T_{\mathrm{RES}}}^{0} d\tau g(\tau)\right]^2 \boldsymbol{\Psi}_0 + k_{\mathrm{INI}}T_{\mathrm{RES}}\overline{g^2}\left\langle n_a \mid \mathrm{E}\right\rangle = k_{\mathrm{INI}}^2 T_{\mathrm{RES}}^2 \bar{g}^2\left\langle n_a \mid \mathrm{E}\right\rangle + k_{\mathrm{INI}}T_{\mathrm{RES}}\overline{g^2}\left\langle n_a \mid \mathrm{E}\right\rangle, \tag{S16}
$$

$$
\left\langle m_a m_b \mid \mathrm{E}\right\rangle = \frac{1}{2}\left[\int_{-T_{\mathrm{RES}}}^{0} d\tau g(\tau)\right]^2 2\mathbf{uK}_{\mathrm{INI},a}\mathbf{K}_{\mathrm{INI},b}\boldsymbol{\Psi}_0 = k_{\mathrm{INI}}^2 T_{\mathrm{RES}}^2 \bar{g}^2\left\langle n_a n_b \mid \mathrm{E}\right\rangle. \tag{S17}
$$

Thus, the variance and covariance are given by:

$$
\begin{aligned}
\sigma_{m|\mathrm{E}}^2 &= \left\langle m_a^2 \mid \mathrm{E}\right\rangle - \left\langle m_a \mid \mathrm{E}\right\rangle^2 = k_{\mathrm{INI}}^2 T_{\mathrm{RES}}^2 \bar{g}^2\left\langle n_a \mid \mathrm{E}\right\rangle + k_{\mathrm{INI}}T_{\mathrm{RES}}\overline{g^2}\left\langle n_a \mid \mathrm{E}\right\rangle - k_{\mathrm{INI}}^2 T_{\mathrm{RES}}^2 \bar{g}^2\left\langle n_a \mid \mathrm{E}\right\rangle^2 \\
&= k_{\mathrm{INI}}^2 T_{\mathrm{RES}}^2 \bar{g}^2 \sigma_{n|\mathrm{E}}^2 + k_{\mathrm{INI}}T_{\mathrm{RES}}\overline{g^2}\left\langle n \mid \mathrm{E}\right\rangle,
\end{aligned}
\tag{S18}
$$

$$
\begin{aligned}
\mathrm{Cov}(m_a, m_b \mid \mathrm{E}) &= \left\langle m_a m_b \mid \mathrm{E}\right\rangle - \left\langle m_a \mid \mathrm{E}\right\rangle\left\langle m_b \mid \mathrm{E}\right\rangle = k_{\mathrm{INI}}^2 T_{\mathrm{RES}}^2 \bar{g}^2\left\langle n_a n_b \mid \mathrm{E}\right\rangle - k_{\mathrm{INI}}^2 T_{\mathrm{RES}}^2 \bar{g}^2\left\langle n_a \mid \mathrm{E}\right\rangle\left\langle n_b \mid \mathrm{E}\right\rangle \\
&= k_{\mathrm{INI}}^2 T_{\mathrm{RES}}^2 \bar{g}^2 \mathrm{Cov}(n_a, n_b \mid \mathrm{E}).
\end{aligned}
\tag{S19}
$$

Therefore, the correlation coefficient between the two gene copies' expression levels satisfies:

$$
\rho_{m_a,m_b|\mathrm{E}} = \frac{\mathrm{Cov}(m_a, m_b \mid \mathrm{E})}{\sigma_{m|\mathrm{E}}^2} = k_{\mathrm{INI}}^2 T_{\mathrm{RES}}^2 \bar{g}^2 \frac{\mathrm{Cov}(n_a, n_b \mid \mathrm{E})}{\sigma_{n|\mathrm{E}}^2}\frac{\sigma_{n|\mathrm{E}}^2}{\sigma_{m|\mathrm{E}}^2} = \rho_{n_a,n_b|\mathrm{E}}\bar{g}^2 k_{\mathrm{INI}}^2 T_{\mathrm{RES}}^2 \frac{\sigma_{n|\mathrm{E}}^2}{\sigma_{m|\mathrm{E}}^2} = \rho_{n_a,n_b|\mathrm{E}}\frac{\eta_{n|\mathrm{E}}^2}{\eta_{m|\mathrm{E}}^2}, \tag{S20}
$$

where $\eta_{n|\mathrm{E}}^2 = \dfrac{\sigma_{n|\mathrm{E}}^2}{\left\langle n \mid \mathrm{E}\right\rangle^2} = \dfrac{1}{\left\langle n \mid \mathrm{E}\right\rangle} - 1$ and $\eta_{m|\mathrm{E}}^2 = \dfrac{\sigma_{m|\mathrm{E}}^2}{\left\langle m \mid \mathrm{E}\right\rangle^2} = \dfrac{1}{\left\langle n \mid \mathrm{E}\right\rangle}\left(1 + \dfrac{\overline{g^2}}{\bar{g}^2 k_{\mathrm{INI}}T_{\mathrm{RES}}}\right) - 1$ are the total noise at the gene state and nascent RNA levels, respectively.

**Supplemental Material**

If the time scale of gene-state transitions is not negligibly small compared to that of nascent RNA production, i.e., $k_{\mathrm{ON}}T_{\mathrm{RES}}, k_{\mathrm{OFF}}T_{\mathrm{RES}} \lesssim 1$ [4,7,31], the second lowest-order terms in **Eqs. (S14)-(S15)** need to be considered:

$$\begin{aligned}\sigma_{m|\mathrm{E}}^2 &\approx k_{\mathrm{INI}}^2 T_{\mathrm{RES}}^2 \bar{g}^2 \sigma_{n|\mathrm{E}}^2 + k_{\mathrm{INI}} T_{\mathrm{RES}} \overline{g^2} \langle n \mid \mathrm{E} \rangle + 2\mathbf{u}\mathbf{K}_{\mathrm{INI},a}\mathbf{K}\mathbf{K}_{\mathrm{INI},a}\mathbf{\Psi}_0 \left[ \int_{-T_{\mathrm{RES}}}^{0} d\tau_1 \int_{-T_{\mathrm{RES}}}^{\tau_1} d\tau_2 g(\tau_1) g(\tau_2)(\tau_1 - \tau_2) \right] \\ &= k_{\mathrm{INI}}^2 T_{\mathrm{RES}}^2 \bar{g}^2 \sigma_{n|\mathrm{E}}^2 + k_{\mathrm{INI}} T_{\mathrm{RES}} \overline{g^2} \langle n \mid \mathrm{E} \rangle - \frac{2k_{\mathrm{INI}}^2 k_{\mathrm{OFF}}(k_{\mathrm{ON}}k_{\mathrm{OFF}} + \gamma k_{\mathrm{ON}}^2)}{k_{\mathrm{OFF}}^2 + 2k_{\mathrm{ON}}k_{\mathrm{OFF}} + \gamma k_{\mathrm{ON}}^2} r_{g,1} \\ &= k_{\mathrm{INI}}^2 T_{\mathrm{RES}}^2 \bar{g}^2 \left[ \sigma_{n|\mathrm{E}}^2 - 2r_{g,1} k_{\mathrm{OFF}} T_{\mathrm{RES}} \langle n \mid \mathrm{E} \rangle \right] + k_{\mathrm{INI}} T_{\mathrm{RES}} \overline{g^2} \langle n \mid \mathrm{E} \rangle, \end{aligned} \tag{S21}$$

$$\begin{aligned}\mathrm{Cov}(m_a, m_b \mid \mathrm{E}) &\approx k_{\mathrm{INI}}^2 T_{\mathrm{RES}}^2 \bar{g}^2 \mathrm{Cov}(n_a, n_b \mid \mathrm{E}) + \mathbf{u}\mathbf{K}_{\mathrm{INI},a}\mathbf{K}^2\mathbf{K}_{\mathrm{INI},b}\mathbf{\Psi}_0 \left[ \int_{-T_{\mathrm{RES}}}^{0} d\tau_1 \int_{-T_{\mathrm{RES}}}^{\tau_1} d\tau_2 g(\tau_1) g(\tau_2)(\tau_1 - \tau_2)^2 \right] \\ &= k_{\mathrm{INI}}^2 T_{\mathrm{RES}}^2 \bar{g}^2 \mathrm{Cov}(n_a, n_b \mid \mathrm{E}) - \frac{(\gamma - 1) k_{\mathrm{INI}}^2 k_{\mathrm{OFF}}^2 k_{\mathrm{ON}}^2}{k_{\mathrm{OFF}}^2 + 2k_{\mathrm{ON}}k_{\mathrm{OFF}} + \gamma k_{\mathrm{ON}}^2} r_{g,2} \\ &= k_{\mathrm{INI}}^2 T_{\mathrm{RES}}^2 \bar{g}^2 \left[ \mathrm{Cov}(n_a, n_b \mid \mathrm{E}) - r_{g,2} \frac{\gamma - 1}{\gamma} (k_{\mathrm{OFF}} T_{\mathrm{RES}})^2 P(n_a = 1, n_b = 1 \mid \mathrm{E}) \right], \end{aligned} \tag{S22}$$

where $r_{g,s} = \dfrac{2\int_{-T_{\mathrm{RES}}}^{0} d\tau_1 \int_{-T_{\mathrm{RES}}}^{\tau_1} d\tau_2 g(\tau_1) g(\tau_2)(\tau_1 - \tau_1)^s}{\bar{g}^2 T_{\mathrm{RES}}^{2+s}}$ are typically small numbers. For example, $r_{g,1} = 0.13$ and $r_{g,2} = 0.0556 \ll 1$ for $g = -\tau / T_{\mathrm{RES}}$.

##### 4.3 Fast gene-state transitions ($k_{\mathrm{ON}}T_{\mathrm{RES}}, k_{\mathrm{OFF}}T_{\mathrm{RES}} \gg 1$)

For $k_{\mathrm{ON}}T_{\mathrm{RES}}, k_{\mathrm{OFF}}T_{\mathrm{RES}} \gg 1$, all nonzero eigenvalues of **K** approaches minus infinity, i.e., . Therefore, $\mathbf{Q}e^{\mathbf{K}_0(\tau_1 - \tau_2)}\mathbf{Q}^{-1} \approx \mathbf{Q} \begin{bmatrix} 1 & 0 & 0 & 0 \\ 0 & 0 & 0 & 0 \\ 0 & 0 & 0 & 0 \\ 0 & 0 & 0 & 0 \end{bmatrix} \mathbf{Q}^{-1} = \left[ \tilde{\mathbf{P}}_0, \tilde{\mathbf{P}}_0, \tilde{\mathbf{P}}_0, \tilde{\mathbf{P}}_0 \right] = \tilde{\mathbf{P}}_0 \mathbf{u}$. This simplifies the second order moments to:

$$\begin{aligned}\langle m_a^2 \mid \mathrm{E} \rangle &= \left[ \int_{-T_{\mathrm{RES}}}^{0} d\tau g(\tau) \right]^2 \mathbf{u}\mathbf{K}_{\mathrm{INI},a}\tilde{\mathbf{P}}_0 \mathbf{u}\mathbf{K}_{\mathrm{INI},a}\mathbf{\Psi}_0 + k_{\mathrm{INI}} T_{\mathrm{RES}} \overline{g^2} \langle n_a \mid \mathrm{E} \rangle \\ &= k_{\mathrm{INI}}^2 T_{\mathrm{RES}}^2 \bar{g}^2 \langle n_a \mid \mathrm{E} \rangle^2 + k_{\mathrm{INI}} T_{\mathrm{RES}} \overline{g^2} \langle n_a \mid \mathrm{E} \rangle, \end{aligned} \tag{S23}$$

$$\langle m_a m_b \mid \mathrm{E}\rangle = \frac{1}{2}\left[\int_{-T_{\mathrm{RES}}}^{0} d\tau g(\tau)\right]^2 2\mathbf{u}\mathbf{K}_{\mathrm{INI},a}\tilde{\mathbf{P}}_0\mathbf{u}\mathbf{K}_{\mathrm{INI},b}\mathbf{\Psi}_0 = k_{\mathrm{INI}}^2 T_{\mathrm{RES}}^2 \bar{g}^2 \langle n_a \mid \mathrm{E}\rangle\langle n_b \mid \mathrm{E}\rangle. \quad \text{(S24)}$$

Thus, the variance and covariance are given by:

$$\sigma_{m|\mathrm{E}}^2 = \langle m_a^2 \mid \mathrm{E}\rangle - \langle m_a \mid \mathrm{E}\rangle^2 = k_{\mathrm{INI}} T_{\mathrm{RES}} \overline{g^2} \langle n_a \mid \mathrm{E}\rangle, \quad \text{(S25)}$$

$$\mathrm{Cov}(m_a, m_b \mid \mathrm{E}) = \langle m_a m_b \mid \mathrm{E}\rangle - \langle m_a \mid \mathrm{E}\rangle\langle m_b \mid \mathrm{E}\rangle = 0. \quad \text{(S26)}$$

I.e., as gene-state transitions become infinitely fast, both the variance and covariance induced by these transitions vanish, leaving only the variance from Poissonian initiation.

#### 5. Deriving the extrinsic fluctuation of nascent RNA

Following the computation of nascent RNA statistics under a given environment, we apply the adiabatic approximation of environmental fluctuations to write the extrinsic variation of nascent RNA level as:

$$\begin{aligned}\sigma_{\mathrm{ext}}^2 = \sigma_{\langle m|\mathrm{E}\rangle}^2 &= \bar{g}^2 \left(\langle k_{\mathrm{INI}}^2 T_{\mathrm{RES}}^2 n_{\mathrm{E}}^2\rangle - \langle k_{\mathrm{INI}} T_{\mathrm{RES}} n_{\mathrm{E}}\rangle^2\right) \\ &= \bar{g}^2 \left(\int k_{\mathrm{INI}}^2 T_{\mathrm{RES}}^2 n_{\mathrm{E}}^2 p(\mathrm{E}) d\mathrm{E} - \left(\int k_{\mathrm{INI}} T_{\mathrm{RES}} n_{\mathrm{E}} p(\mathrm{E}) d\mathrm{E}\right)^2\right). \quad \text{(S27)}\end{aligned}$$

Here, $k_{\mathrm{INI}}$, $T_{\mathrm{RES}}$, and $n_{\mathrm{E}}$ represent quantities under a given environment with a given set of kinetic parameters ($\mathrm{E} = [k_{\mathrm{ON}}, k_{\mathrm{OFF}}, k_{\mathrm{INI}}, T_{\mathrm{RES}}, \gamma]$), while $\langle \cdot \rangle$ denotes averaging over fluctuating environments with a probability distribution $p(\mathrm{E})$. Since $n_{\mathrm{E}} \equiv \langle n \mid \mathrm{E}\rangle$ only depends on $k_{\mathrm{ON}} / k_{\mathrm{OFF}}$ and $\gamma$, the arguments of $p(\mathrm{E})$ can be regrouped into three variables, i.e., $k_{\mathrm{INI}} T_{\mathrm{RES}}$, $k_{\mathrm{ON}} / k_{\mathrm{OFF}}$, and $\gamma$. Assuming these variables are independently controlled by different external factors, the extrinsic noise can be further decomposed into:

$$\begin{aligned}\sigma_{\mathrm{ext}}^2 &= \bar{g}^2 \left(\langle k_{\mathrm{INI}}^2 T_{\mathrm{RES}}^2\rangle\langle n_{\mathrm{E}}^2\rangle - \langle k_{\mathrm{INI}} T_{\mathrm{RES}}\rangle^2 \langle n_{\mathrm{E}}\rangle^2\right) \\ &= \bar{g}^2 \left(\langle k_{\mathrm{INI}}^2 T_{\mathrm{RES}}^2\rangle\langle n_{\mathrm{E}}^2\rangle - \langle k_{\mathrm{INI}}^2 T_{\mathrm{RES}}^2\rangle\langle n_{\mathrm{E}}\rangle^2 + \langle k_{\mathrm{INI}}^2 T_{\mathrm{RES}}^2\rangle\langle n_{\mathrm{E}}\rangle^2 - \langle k_{\mathrm{INI}} T_{\mathrm{RES}}\rangle^2 \langle n_{\mathrm{E}}\rangle^2\right) \\ &= \bar{g}^2 \langle k_{\mathrm{INI}}^2 T_{\mathrm{RES}}^2\rangle \sigma_{\langle n|\mathrm{E}\rangle}^2 + \bar{g}^2 \langle n\rangle^2 \sigma_{\langle k_{\mathrm{INI}} T_{\mathrm{RES}}|\mathrm{E}\rangle}^2, \quad \text{(S28)}\end{aligned}$$

where the two terms represent the variances arising from extrinsic fluctuations in ON-state probability and transcription initiation (or residence time), respectively. According to previous studies, $k_{\mathrm{ON}}$ and $k_{\mathrm{OFF}}$ are typically more variable than $k_{\mathrm{INI}}$ [23,24]. Therefore, the second term may be neglected, i.e.,

$$\sigma_{\mathrm{ext}}^2 \approx \bar{g}^2 k_{\mathrm{INI}}^2 T_{\mathrm{RES}}^2 \sigma_{\langle n|\mathrm{E}\rangle}^2. \tag{S29}$$

#### 6. Deriving the coupled intrinsic fluctuation in noisy environments

##### 6.1 Zeroth-order results

To integrate intrinsic coupling with environmental changes, we start by assuming slow gene-state transitions ($k_{\mathrm{ON}} T_{\mathrm{RES}},\ k_{\mathrm{OFF}} T_{\mathrm{RES}} \ll 1$). In this limiting case, combining **Eqs. (S19)** and **(S4)** leads to:

$$\begin{aligned}\mathrm{Cov}(m_a, m_b \mid \mathrm{E}) &= k_{\mathrm{INI}}^2 T_{\mathrm{RES}}^2 \bar{g}^2 \mathrm{Cov}(n_a, n_b \mid \mathrm{E}) \\ &= \bar{g}^2 k_{\mathrm{INI}}^2 T_{\mathrm{RES}}^2 \underbrace{(\gamma-1)\left(\frac{(1-n_{\mathrm{E}})\left(2n_{\mathrm{E}}-1+\sqrt{4(\gamma-1)n_{\mathrm{E}}(1-n_{\mathrm{E}})+1}\right)}{2\gamma(1-n_{\mathrm{E}})+2n_{\mathrm{E}}-1+\sqrt{4(\gamma-1)n_{\mathrm{E}}(1-n_{\mathrm{E}})+1}}\right)^2}_{f(n_{\mathrm{E}},\gamma)}.\end{aligned} \tag{S30}$$

Notably, $f$ is symmetric with respect to $n_{\mathrm{E}} = 0.5$, and $f(1,\gamma) = f(0,\gamma) = f(n_{\mathrm{E}},1) = 0$. The coupled intrinsic fluctuation in changing environments can be derived by taking a series expansion of $f$, i.e.:

$$\begin{aligned}\sigma_{\mathrm{int,cp}}^2 &= \langle \mathrm{Cov}(m_a, m_b \mid \mathrm{E}) \rangle = \bar{g}^2 k_{\mathrm{INI}}^2 T_{\mathrm{RES}}^2 \left\langle f(\langle n \mid \mathrm{E} \rangle, \gamma) \right\rangle \\ &\approx \bar{g}^2 k_{\mathrm{INI}}^2 T_{\mathrm{RES}}^2 \left\langle f(\langle n \rangle, \gamma) + \left.\frac{\partial f}{\partial n}\right|_{\langle n \rangle} (\langle n \mid \mathrm{E} \rangle - \langle n \rangle) + \frac{1}{2}\left.\frac{\partial^2 f}{\partial n^2}\right|_{\langle n \rangle} (\langle n \mid \mathrm{E} \rangle - \langle n \rangle)^2 + \cdots \right\rangle \\ &\approx \bar{g}^2 k_{\mathrm{INI}}^2 T_{\mathrm{RES}}^2 \left[ f(\langle n \rangle, \gamma) + \frac{1}{2}\left.\frac{\partial^2 f}{\partial n^2}\right|_{\langle n \rangle} \sigma_{\langle n|\mathrm{E}\rangle}^2 \right].\end{aligned} \tag{S31}$$

Here, similar to the last section, we have assumed that $k_{\mathrm{ON}}$ and $k_{\mathrm{OFF}}$ exhibit greater variability than other parameters (such as $k_{\mathrm{INI}}$ and $\gamma$) in fluctuating environments[23,24]. Combining **Eqs. (S29)** and **(S31)** leads to the expression of total covariance:

$$\mathrm{Cov}(m_a, m_b) \approx \bar{g}^2 k_{\mathrm{INI}}^2 T_{\mathrm{RES}}^2 \left[ f(\langle n \rangle, \gamma) + \left( \frac{1}{2}\left.\frac{\partial^2 f}{\partial n^2}\right|_{\langle n \rangle} + 1 \right) \sigma_{\langle n|\mathrm{E}\rangle}^2 \right]. \tag{S32}$$

#### 6.2 Correction for $k_{\mathrm{ON}}T_{\mathrm{RES}}, k_{\mathrm{OFF}}T_{\mathrm{RES}} \lesssim 1$

Note that **Eqs. (S30)-(S31)** are derived in the limit of slow gene-state transitions ($k_{\mathrm{ON}}T_{\mathrm{RES}}, k_{\mathrm{OFF}}T_{\mathrm{RES}} \ll 1$). Outside this limit (i.e., $k_{\mathrm{ON}}T_{\mathrm{RES}}, k_{\mathrm{OFF}}T_{\mathrm{RES}} \lesssim 1$), the covariance between $m_a$ and $m_b$ under a given environmental condition also depends on $k_{\mathrm{OFF}}$, i.e.:

$$\begin{aligned}\mathrm{Cov}(m_a,m_b\,|\,\mathrm{E}) &\approx \bar{g}^2 k_{\mathrm{INI}}^2 T_{\mathrm{RES}}^2\left[f(\langle n\,|\,\mathrm{E}\rangle,\gamma)-r_{g,2}\frac{\gamma-1}{\gamma}(k_{\mathrm{OFF}}T_{\mathrm{RES}})^2 P(n_a=1,n_b=1)\right] \\ &\approx \bar{g}^2 k_{\mathrm{INI}}^2 T_{\mathrm{RES}}^2\left[\left(1-\frac{\gamma-1}{\gamma}r_{g,2}k_{\mathrm{OFF}}^2T_{\mathrm{RES}}^2\right)f(\langle n\,|\,\mathrm{E}\rangle,\gamma)-\frac{\gamma-1}{\gamma}r_{g,2}k_{\mathrm{OFF}}^2T_{\mathrm{RES}}^2\langle n\,|\,\mathrm{E}\rangle^2\right].\end{aligned} \tag{S33}$$

If environmental fluctuations primarily affect $k_{\mathrm{ON}}$ ($k_{\mathrm{OFF}}$ stays relatively stable), **Eq. (S31)** is corrected to:

$$\begin{aligned}\sigma_{\mathrm{int,cp}}^2 = \langle \mathrm{Cov}(m_a,m_b\,|\,\mathrm{E})\rangle &\approx \bar{g}^2 k_{\mathrm{INI}}^2 T_{\mathrm{RES}}^2\left[\left(1-\frac{\gamma-1}{\gamma}r_{g,2}k_{\mathrm{OFF}}^2T_{\mathrm{RES}}^2\right)f(\langle n\rangle,\gamma)-\frac{\gamma-1}{\gamma}r_{g,2}k_{\mathrm{OFF}}^2T_{\mathrm{RES}}^2\langle n\rangle^2\right] \\ &+\bar{g}^2 k_{\mathrm{INI}}^2 T_{\mathrm{RES}}^2\left[\frac{1}{2}\left(1-\frac{\gamma-1}{\gamma}r_{g,2}k_{\mathrm{OFF}}^2T_{\mathrm{RES}}^2\right)\left.\frac{\partial^2 f}{\partial n^2}\right|_{\langle n\rangle}-\frac{\gamma-1}{\gamma}r_{g,2}k_{\mathrm{OFF}}^2T_{\mathrm{RES}}^2\right]\sigma_{\langle n|\mathrm{E}\rangle}^2.\end{aligned} \tag{S34}$$

Therefore, the total covariance is written as:

$$\begin{aligned}\mathrm{Cov}(m_a,m_b) &\approx \bar{g}^2 k_{\mathrm{INI}}^2 T_{\mathrm{RES}}^2\left[\left(1-\frac{\gamma-1}{\gamma}r_{g,2}k_{\mathrm{OFF}}^2T_{\mathrm{RES}}^2\right)f(\langle n\rangle,\gamma)-\frac{\gamma-1}{\gamma}r_{g,2}k_{\mathrm{OFF}}^2T_{\mathrm{RES}}^2\langle n\rangle^2\right] \\ &+\bar{g}^2 k_{\mathrm{INI}}^2 T_{\mathrm{RES}}^2\left(1-\frac{\gamma-1}{\gamma}r_{g,2}k_{\mathrm{OFF}}^2T_{\mathrm{RES}}^2\right)\left(\frac{1}{2}\left.\frac{\partial^2 f}{\partial n^2}\right|_{\langle n\rangle}+1\right)\sigma_{\langle n|\mathrm{E}\rangle}^2.\end{aligned} \tag{S35}$$

Interestingly, compared to **Eq. (S32)**, the $\sigma_{\langle n|\mathrm{E}\rangle}^2$ term in **Eq. (S35)** differs only by an additional coefficient $\left(1-\frac{\gamma-1}{\gamma}r_{g,2}k_{\mathrm{OFF}}^2T_{\mathrm{RES}}^2\right)$, which is typically very close to one as $r_{g,2} \ll 1$. Therefore, the proposed framework for estimating $\gamma$ and decomposing correlated noise holds for $k_{\mathrm{OFF}}T_{\mathrm{RES}} \lesssim 1$. Alternatively, we describe a different $\gamma$-estimation method in **Section 9** to explicitly account for this correction.

**Supplemental Material**

**7. Deriving the total correlated fluctuation for $\gamma = 0, 1, \text{and} \infty$**

While determining the dependence of total covariance on $\langle n \rangle$ and $\sigma^2_{\langle n|\mathrm{E}\rangle}$ for arbitrary relies on the Taylor expansion of $f$, simple results are available for the following $\gamma$ values:

1. For $\gamma = 1$ (independent gene pair), $f = 0$, which leads to

$$\mathrm{Cov}(m_a, m_b) = \langle \mathrm{Cov}(m_a, m_b \mid \mathrm{E}) \rangle + \sigma^2_{\langle m|\mathrm{E}\rangle} = \sigma^2_{\langle m|\mathrm{E}\rangle} = \bar{g}^2 k^2_{\mathrm{INI}} T^2_{\mathrm{RES}} \sigma^2_{\langle n|\mathrm{E}\rangle}. \tag{S36}$$

Therefore, $\mathrm{Cov}(m_a, m_b)$ is a function of $\sigma^2_{\langle n|\mathrm{E}\rangle}$, but independent of $\langle n \rangle$.

2. For $\gamma \to \infty$ (fully coupled gene pair), we have:

$$\lim_{\gamma\to\infty} f(n_{\mathrm{E}}, \gamma) = \lim_{\gamma\to\infty} (\gamma - 1)\left( \frac{(1-n_{\mathrm{E}})\left(2n_{\mathrm{E}} - 1 + \sqrt{4(\gamma-1)n_{\mathrm{E}}(1-n_{\mathrm{E}})+1}\right)}{2\gamma(1-n_{\mathrm{E}}) + 2n_{\mathrm{E}} - 1 + \sqrt{4(\gamma-1)n_{\mathrm{E}}(1-n_{\mathrm{E}})+1}} \right)^2 = n_{\mathrm{E}}(1-n_{\mathrm{E}}) = \sigma^2_{n|\mathrm{E}}. \tag{S37}$$

Hence, the total covariance simplifies to:

$$\mathrm{Cov}(m_a, m_b) = \langle \mathrm{Cov}(m_a, m_b \mid \mathrm{E}) \rangle + \sigma^2_{\langle m|\mathrm{E}\rangle} = \bar{g}^2 k^2_{\mathrm{INI}} T^2_{\mathrm{RES}} \left[ \left\langle \sigma^2_{n|\mathrm{E}} \right\rangle + \sigma^2_{\langle n|\mathrm{E}\rangle} \right] = \bar{g}^2 k^2_{\mathrm{INI}} T^2_{\mathrm{RES}} \langle n \rangle \left(1 - \langle n \rangle\right), \tag{S38}$$

which depends exclusively on $\langle n \rangle$.

3. For $\gamma \to 0$ (fully anti-coupled gene pair), we have:

$$\lim_{\gamma\to 0} f(n_{\mathrm{E}}, \gamma) = \lim_{\gamma\to 0} (\gamma - 1)\left( \frac{(1-n_{\mathrm{E}})\left(2n_{\mathrm{E}} - 1 + \sqrt{4(\gamma-1)n_{\mathrm{E}}(1-n_{\mathrm{E}})+1}\right)}{2\gamma(1-n_{\mathrm{E}}) + 2n_{\mathrm{E}} - 1 + \sqrt{4(\gamma-1)n_{\mathrm{E}}(1-n_{\mathrm{E}})+1}} \right)^2 = -(1-n_{\mathrm{E}})^2. \tag{S39}$$

Hence, the total covariance becomes:

$$\begin{aligned}\mathrm{Cov}(m_a, m_b) &= \langle \mathrm{Cov}(m_a, m_b \mid \mathrm{E}) \rangle + \sigma^2_{\langle m|\mathrm{E}\rangle} = -\bar{g}^2 k^2_{\mathrm{INI}} T^2_{\mathrm{RES}} \left\langle \left(1 - \langle n \mid \mathrm{E} \rangle\right)^2 \right\rangle + \bar{g}^2 k^2_{\mathrm{INI}} T^2_{\mathrm{RES}} \sigma^2_{\langle n|\mathrm{E}\rangle} \\ &= \bar{g}^2 k^2_{\mathrm{INI}} T^2_{\mathrm{RES}} \left[ \left\langle \sigma^2_{n|\mathrm{E}} \right\rangle + \langle n \rangle - 1 + \sigma^2_{\langle n|\mathrm{E}\rangle} \right] = -\bar{g}^2 k^2_{\mathrm{INI}} T^2_{\mathrm{RES}} \left(1 - \langle n \rangle\right)^2, \end{aligned} \tag{S40}$$

which, similar to Case 2, is only dependent on $\langle n \rangle$, but not on $\sigma^2_{\langle n|\mathrm{E}\rangle}$.

**Supplemental Material**

#### 8. Deriving the total transcriptional fluctuation for a single gene

##### 8.1 Zeroth-order results

In the limit of slow gene-state transitions ($k_{\mathrm{ON}}T_{\mathrm{RES}},\ k_{\mathrm{OFF}}T_{\mathrm{RES}} \ll 1$), the total variance for each gene copy under fluctuating environments can be derived from **Eqs. (S18)** and **(S29)** as:

$$\begin{aligned}\sigma_m^2 = \left\langle \sigma_{m|\mathrm{E}}^2 \right\rangle + \sigma_{\langle m|\mathrm{E}\rangle}^2 &\approx \bar{g}^2 k_{\mathrm{INI}}^2 T_{\mathrm{RES}}^2 \left\langle \sigma_{n|\mathrm{E}}^2 \right\rangle + \overline{g^2} k_{\mathrm{INI}} T_{\mathrm{RES}} \left\langle \langle n \,|\, \mathrm{E} \rangle \right\rangle + \bar{g}^2 k_{\mathrm{INI}}^2 T_{\mathrm{RES}}^2 \sigma_{\langle n|\mathrm{E}\rangle}^2 \\ &= \bar{g}^2 k_{\mathrm{INI}}^2 T_{\mathrm{RES}}^2 \left[ \left\langle \sigma_{n|\mathrm{E}}^2 \right\rangle + \sigma_{\langle n|\mathrm{E}\rangle}^2 \right] + \overline{g^2} k_{\mathrm{INI}} T_{\mathrm{RES}} \langle n \rangle \\ &= \bar{g}^2 k_{\mathrm{INI}}^2 T_{\mathrm{RES}}^2 \sigma_n^2 + \overline{g^2} k_{\mathrm{INI}} T_{\mathrm{RES}} \langle n \rangle. \end{aligned} \tag{S41}$$

Here, similar to previous sections, we assume that variations in $k_{\mathrm{ON}}$ and $k_{\mathrm{OFF}}$ are the primary contributors to extrinsic fluctuations[23,24]. Therefore, the total noise for a single gene is given by:

$$\begin{aligned}\eta_m^2 = \frac{\sigma_m^2}{\langle m \rangle^2} &\approx \frac{\bar{g}^2 k_{\mathrm{INI}}^2 T_{\mathrm{RES}}^2 \sigma_n^2 + \overline{g^2} k_{\mathrm{INI}} T_{\mathrm{RES}} \langle n \rangle}{\left( \bar{g} k_{\mathrm{INI}} T_{\mathrm{RES}} \langle n \rangle \right)^2} = \frac{\sigma_n^2}{\langle n \rangle^2} + \frac{\overline{g^2}}{\bar{g}^2 k_{\mathrm{INI}} T_{\mathrm{RES}} \langle n \rangle} \\ &= \frac{1}{\langle n \rangle} \left( 1 + \frac{\overline{g^2}}{\bar{g}^2 k_{\mathrm{INI}} T_{\mathrm{RES}}} \right) - 1 \approx \frac{1}{\langle n \rangle} - 1. \end{aligned} \tag{S42}$$

The last approximation requires $k_{\mathrm{INI}} T_{\mathrm{RES}} \gg 1$, which holds for many biological systems[5–7,14,31,32].

##### 8.2 Correction for $k_{\mathrm{ON}}T_{\mathrm{RES}}, k_{\mathrm{OFF}}T_{\mathrm{RES}} \lesssim 1$

If gene-state transitions are not extremely slow (i.e., $k_{\mathrm{ON}}T_{\mathrm{RES}},\ k_{\mathrm{OFF}}T_{\mathrm{RES}} \lesssim 1$), **Eq. (S41)** needs to be corrected by adding an extra $k_{\mathrm{OFF}}$-dependent term from **Eq. (S21)**, i.e.:

$$\sigma_m^2 \approx \bar{g}^2 k_{\mathrm{INI}}^2 T_{\mathrm{RES}}^2 \sigma_n^2 + \overline{g^2} k_{\mathrm{INI}} T_{\mathrm{RES}} \langle n \rangle - 2 r_{g,1} k_{\mathrm{OFF}} T_{\mathrm{RES}} \bar{g}^2 k_{\mathrm{INI}}^2 T_{\mathrm{RES}}^2 \langle n \rangle, \tag{S43}$$

where we have assumed that environmental fluctuations primarily affect $k_{\mathrm{ON}}$ ($k_{\mathrm{OFF}}$ stays relatively stable), as in previous sections. Therefore, **Eq. (S31)** is modified to:

$$\eta_m^2 \approx \frac{1}{\langle n \rangle} \left( 1 - 2 r_{g,1} k_{\mathrm{OFF}} T_{\mathrm{RES}} + \frac{\overline{g^2}}{\bar{g}^2 k_{\mathrm{INI}} T_{\mathrm{RES}}} \right) - 1 \approx \frac{1 - 2 r_{g,1} k_{\mathrm{OFF}} T_{\mathrm{RES}}}{\langle n \rangle} - 1. \tag{S44}$$

Since $r_{g,2} < 1$ is a small factor, this correction is typically negligible. Alternatively, this correction may be incorporated in a different method for $\gamma$-estimation, as described in **Section 9**.

#### 9. Alternative methods for determining $\gamma$

Two alternative methods may be employed as replacements for Steps 2–3 in the noise decomposition framework outlined in the main text to determine $\gamma$:

(1) By measuring $\frac{\partial \mathrm{Cov}(m_a, m_b)}{\partial \sigma^2_{\langle m|\mathrm{E}\rangle}}$ from two values of $\langle n \rangle$ ($\langle n_1 \rangle$ and $\langle n_2 \rangle$), we can compute their ratio:

$$\left.\frac{\partial \mathrm{Cov}(m_a, m_b)}{\partial \sigma^2_{\langle m|\mathrm{E}\rangle}}\right|_{\langle n_2 \rangle} \Bigg/ \left.\frac{\partial \mathrm{Cov}(m_a, m_b)}{\partial \sigma^2_{\langle m|\mathrm{E}\rangle}}\right|_{\langle n_1 \rangle} = \left(\frac{1}{2}\left.\frac{\partial^2 f}{\partial n^2}\right|_{\langle n_2 \rangle} + 1\right) \Bigg/ \left(\frac{1}{2}\left.\frac{\partial^2 f}{\partial n^2}\right|_{\langle n_1 \rangle} + 1\right), \tag{S45}$$

which is a monotonic function of $\gamma$. This allows us to uniquely determine $\gamma$ (**Fig. S2(b)**). Notably, **Eq. (S45)** holds true even when accounting for the second-order term in $\mathrm{Cov}(m_a, m_b \mid \mathrm{E})$. Therefore, it is theoretically more precise for determining $\gamma$ beyond the limit of slow gene-state transitions. Here, by comparing results from two sets of $\langle n_1 \rangle$ and $\langle n_2 \rangle$, we can further determine whether a correction for $\langle n \rangle$ is necessary, as mention in **Section 8**. If needed, this correction (a scaling factor for $\langle n \rangle$) can be solved alongside $\gamma$ from two sets of **Eq. (S45)**.

(2) If the absolute value of $\sigma^2_{\langle m|\mathrm{E}\rangle}$ can be measured (e.g. from a control system without coupling), we can extrapolate the linear relationship between $\mathrm{Cov}(m_a, m_b)$ and $\sigma^2_{\langle m|\mathrm{E}\rangle}$ to $\sigma^2_{\langle m|\mathrm{E}\rangle} = 0$, which yields:

$$\mathrm{Cov}(m_a, m_b)\big|_{\sigma^2_{\langle n|\mathrm{E}\rangle}=0} \approx \bar{g}^2 k^2_{\mathrm{INI}} f(\langle n \rangle, \gamma). \tag{S46}$$

With $\langle n \rangle$ and $\bar{g}^2 k_{\text{INI}}^2$ estimated from Step 1 of the noise decomposition framework, a unique solution for $\gamma$ can be determined (**Fig. S2(c)**). This method only requires measurements at one value of $\langle n \rangle$.

#### 10. Separating extrinsic fluctuation and coupled intrinsic fluctuation

With given $\gamma$ and $\langle n \rangle$, we can apply **Eqs. (12)** and **(13)** to decompose total covariance into:

$$\begin{cases} \sigma_{\text{int,cp}}^2 = \bar{g}^2 k_{\text{INI}}^2 T_{\text{RES}}^2 f(\langle n \rangle, \gamma) + \dfrac{\partial_n^2 f\big|_{\langle n \rangle}}{\partial_n^2 f\big|_{\langle n \rangle} + 2}\left[\text{Cov}(m_a, m_b) - \bar{g}^2 k_{\text{INI}}^2 T_{\text{RES}}^2 f(\langle n \rangle, \gamma)\right] \\ \sigma_{\text{ext}}^2 = \dfrac{2}{\partial_n^2 f\big|_{\langle n \rangle} + 2}\left[\text{Cov}(m_a, m_b) - \bar{g}^2 k_{\text{INI}}^2 T_{\text{RES}}^2 f(\langle n \rangle, \gamma)\right] \end{cases}. \tag{S47}$$

Notably, **Eq. (S47)** does not require prior knowledge of the absolute value of $\sigma_{\langle n|\text{E}\rangle}^2$.

#### 11. Experimental data extraction

We validate our noise decomposition framework using single-cell imaging data of the *hb* gene in developing *Drosophila melanogaster* embryos during nuclear cycles (nc) 11–14. Previous studies have reported that *hb* exhibits bursty transcription kinetics, typically modeled as a two-state telegraph process satisfying $k_{\text{ON}} T_{\text{RES}}$, $k_{\text{OFF}} T_{\text{RES}} \lesssim 1$[6,7,18]. In this work, we use data from a previous experimental study[7]:

**(1) Fixed-embryo imaging of nascent RNA at individual *hb* alleles during late mitotic interphase of each nuclear cycle**. At this stage, each *hb* gene copy has been replicated into two. Nascent RNA from each *hb* allele was labeled via smFISH using probes targeting the coding sequence (CDS) region of the *hb* sequence, without distinguishing RNA from different promoters. Embryos were imaged using high-resolution confocal microscopy to ensure the distinction between signals from closely located sister alleles. In this study, we use six embryos at nc12.

**Supplemental Material**

**(2) Fixed-embryo imaging of nascent RNAs produced from two alternative promoters (P1 and P2) on individual *hb* alleles**. Here, nascent RNA from each promoter was labeled and imaged via smFISH using probes targeting their unique sequence regions (5'UTR of P1-mRNA and 3'UTR of P2-mRNA). This enabled promoter-specific imaging of nascent transcription using high-resolution confocal microscopy. In this study, we use five embryos at nc12–13.

For each imaged embryo, nascent RNA signals within individual nuclei were identified as bright fluorescent spots, with their intensities quantified in absolute molecule numbers using custom MATLAB scripts, following previous literature[6,7]. The spot identification algorithm was updated to improve the distinction between closely located nascent RNA signals from sister alleles.

For **imaging data (1)**, Each *hb* nascent RNA spot was considered an active *hb* allele. Given that each nucleus contains four copies of the *hb* gene during late mitotic interphase, the number of unmarked silent *hb* alleles in each nucleus was estimated based on the number of marked alleles. For nuclei with more than two active *hb* alleles, sister and non-sister *hb* alleles were distinguished using the *k*-means clustering method. For nuclei with two active alleles, sister and non-sister *hb* alleles were distinguished using proximity analysis with a distance threshold of 0.71 μm.

For **imaging data (2)**, nascent RNA spots of the *hb* P1 and P2 promoters were paired based on proximity analysis with a distance threshold of 0.71 μm. Unpaired spots were considered as *hb* loci with only one promoter active. The number of unidentified *hb* loci (those with both promoters silent) was estimated for each nucleus based on the number of identified loci.

#### 12. Model extension for non-identical gene copies

Our model can be generalized to describe the coupling between non-identical gene copies ($i = a$, $b$), each with distinct kinetic parameters ($k_{\mathrm{ON},i}$, $k_{\mathrm{OFF},i}$, $k_{\mathrm{INI},i}$, and $g_i(\tau)$). Here, the potential difference in two genes' $T_{\mathrm{RES}}$ can be addressed by adopting the longer $T_{\mathrm{RES}}$ for both gene copies, while adjusting $g_i(\tau)$ to reflect their difference in RNA residence time (see **Table S1**). The coupling between the two genes is assumed to be reciprocal, with a single coupling strength $\gamma$, i.e., $k_{\mathrm{ON},a} \xrightarrow{n_b=1} \gamma k_{\mathrm{ON},a}$ and $k_{\mathrm{ON},b} \xrightarrow{n_a=1} \gamma k_{\mathrm{ON},b}$.

In this generalized model, **Eq. (3)** remains valid under the following redefinition of its terms, i.e.,

$$\mathbf{K} = \begin{bmatrix} -k_{\mathrm{ON},a} - k_{\mathrm{ON},b} & k_{\mathrm{OFF},a} & k_{\mathrm{OFF},b} & 0 \\ k_{\mathrm{ON},a} & -\gamma k_{\mathrm{ON},b} - k_{\mathrm{OFF},a} & 0 & k_{\mathrm{OFF},b} \\ k_{\mathrm{ON},b} & 0 & -\gamma k_{\mathrm{ON},a} - k_{\mathrm{OFF},b} & k_{\mathrm{OFF},a} \\ 0 & \gamma k_{\mathrm{ON},b} & \gamma k_{\mathrm{ON},a} & -k_{\mathrm{OFF},a} - k_{\mathrm{OFF},b} \end{bmatrix}, \quad \mathbf{K}_{\mathrm{INI},a} = \begin{bmatrix} 0 & 0 & 0 & 0 \\ 0 & k_{\mathrm{INI},a} & 0 & 0 \\ 0 & 0 & 0 & 0 \\ 0 & 0 & 0 & k_{\mathrm{INI},a} \end{bmatrix},$$

and $\mathbf{K}_{\mathrm{INI},b} = \begin{bmatrix} 0 & 0 & 0 & 0 \\ 0 & 0 & 0 & 0 \\ 0 & 0 & k_{\mathrm{INI},b} & 0 \\ 0 & 0 & 0 & k_{\mathrm{INI},b} \end{bmatrix}$. The steady-state marginal distribution of gene states satisfies

$$\mathbf{P}_{n_a,n_b|\mathrm{E}} = \frac{1}{1 + \dfrac{k_{\mathrm{ON},a}}{k_{\mathrm{OFF},a}} + \dfrac{k_{\mathrm{ON},b}}{k_{\mathrm{OFF},b}} + \gamma \dfrac{k_{\mathrm{ON},a}}{k_{\mathrm{OFF},a}} \dfrac{k_{\mathrm{ON},b}}{k_{\mathrm{OFF},b}}} \begin{bmatrix} 1 \\ \dfrac{k_{\mathrm{ON},a}}{k_{\mathrm{OFF},a}} \\ \dfrac{k_{\mathrm{ON},b}}{k_{\mathrm{OFF},b}} \\ \gamma \dfrac{k_{\mathrm{ON},a}}{k_{\mathrm{OFF},a}} \dfrac{k_{\mathrm{ON},b}}{k_{\mathrm{OFF},b}} \end{bmatrix}, \tag{S48}$$

which yields:

$$P(n_a = 1 \mid \mathrm{E}) = \langle n_a \mid \mathrm{E} \rangle = \frac{\dfrac{k_{\mathrm{ON},a}}{k_{\mathrm{OFF},a}} + \gamma \dfrac{k_{\mathrm{ON},a}}{k_{\mathrm{OFF},a}} \dfrac{k_{\mathrm{ON},b}}{k_{\mathrm{OFF},b}}}{1 + \dfrac{k_{\mathrm{ON},a}}{k_{\mathrm{OFF},a}} + \dfrac{k_{\mathrm{ON},b}}{k_{\mathrm{OFF},b}} + \gamma \dfrac{k_{\mathrm{ON},a}}{k_{\mathrm{OFF},a}} \dfrac{k_{\mathrm{ON},b}}{k_{\mathrm{OFF},b}}}, \tag{S49}$$

**Supplemental Material**

$$P(n_b = 1 | \mathrm{E}) = \langle n_b | \mathrm{E} \rangle = \frac{\dfrac{k_{\mathrm{ON},b}}{k_{\mathrm{OFF},b}} + \gamma \dfrac{k_{\mathrm{ON},a}}{k_{\mathrm{OFF},a}} \dfrac{k_{\mathrm{ON},b}}{k_{\mathrm{OFF},b}}}{1 + \dfrac{k_{\mathrm{ON},a}}{k_{\mathrm{OFF},a}} + \dfrac{k_{\mathrm{ON},b}}{k_{\mathrm{OFF},b}} + \gamma \dfrac{k_{\mathrm{ON},a}}{k_{\mathrm{OFF},a}} \dfrac{k_{\mathrm{ON},b}}{k_{\mathrm{OFF},b}}}, \tag{S50}$$

$$\mathrm{Cov}(n_a, n_b | \mathrm{E}) = (\gamma - 1) \frac{\dfrac{k_{\mathrm{ON},a}}{k_{\mathrm{OFF},a}} \dfrac{k_{\mathrm{ON},b}}{k_{\mathrm{OFF},b}}}{\left(1 + \dfrac{k_{\mathrm{ON},a}}{k_{\mathrm{OFF},a}} + \dfrac{k_{\mathrm{ON},b}}{k_{\mathrm{OFF},b}} + \gamma \dfrac{k_{\mathrm{ON},a}}{k_{\mathrm{OFF},a}} \dfrac{k_{\mathrm{ON},b}}{k_{\mathrm{OFF},b}}\right)^2}. \tag{S51}$$

These expressions all depend on $\dfrac{k_{\mathrm{ON},a}}{k_{\mathrm{OFF},a}}$, $\dfrac{k_{\mathrm{ON},b}}{k_{\mathrm{OFF},b}}$, and $\gamma$, which, in principle, allow the covariance and correlation coefficient to be written as functions of $\langle n_a | \mathrm{E} \rangle$, $\langle n_b | \mathrm{E} \rangle$, and $\gamma$. Here we consider a specific case where $\dfrac{k_{\mathrm{ON},b}}{k_{\mathrm{OFF},b}}$ is always proportional to $\dfrac{k_{\mathrm{ON},a}}{k_{\mathrm{OFF},a}}$ with a ratio of $\beta$. This assumption makes sense for gene (or promoter) pairs activated by the same enhancer, with $\beta$ representing the difference in the activatability between two genes (or promoters). Based on this assumption, the covariance between $n_a$ and $n_b$ can be written as a function of $\langle n_a | \mathrm{E} \rangle$ (abbreviated as $n_{a\mathrm{E}}$) and $\gamma$ (**Fig. S3(b)**):

$$\begin{aligned} \mathrm{Cov}(n_a, n_b | \mathrm{E}) &= f_\beta(n_{a\mathrm{E}}, \gamma) \\ &= \frac{(\gamma - 1)}{\beta} \left[ \frac{(1 - n_{a\mathrm{E}})\left((1+\beta) n_{a\mathrm{E}} - 1 + \sqrt{(n_{a\mathrm{E}} + \beta n_{a\mathrm{E}} - 1)^2 + 4\beta\gamma n_{a\mathrm{E}}(1 - n_{a\mathrm{E}})}\right)}{2\gamma(1 - n_{a\mathrm{E}}) + (1+\beta) n_{a\mathrm{E}} - 1 + \sqrt{(n_{a\mathrm{E}} + \beta n_{a\mathrm{E}} - 1)^2 + 4\beta\gamma n_{a\mathrm{E}}(1 - n_{a\mathrm{E}})}} \right]^2. \end{aligned} \tag{S52}$$

For a given $\gamma$, the extreme value of covariance appears at $k_{\mathrm{ON}} / k_{\mathrm{OFF}} = (\beta\gamma)^{-1/2}$, which corresponds to $\langle n_a | \mathrm{E} \rangle = \dfrac{\beta^{1/2} + \gamma^{1/2}}{\beta^{1/2} + \beta^{-1/2} + 2\gamma^{1/2}}$.

Similarly, $\langle n_b | \mathrm{E} \rangle$ can be written as a function of $n_{a\mathrm{E}}$ and $\gamma$:

$$\langle n_b \mid \mathrm{E} \rangle = h_\beta(n_{a\mathrm{E}}, \gamma) = \frac{(\beta + n_{a\mathrm{E}} - 1)x + n_{a\mathrm{E}}}{1 + \beta x}, \tag{S53}$$

where $x = \dfrac{[(\beta+1)n_{a\mathrm{E}} - 1] + \sqrt{[(\beta+1)n_{a\mathrm{E}} - 1]^2 + 4\beta\gamma n_{a\mathrm{E}}(1 - n_{a\mathrm{E}})}}{2\beta\gamma(1 - n_{a\mathrm{E}})}$.

Under a given environmental condition, the mean, variance, and covariance at the nascent RNA level follow equations similar to those presented in the main text and earlier sections of the Supplemental Material. Specifically:

$$\langle m_a \mid \mathrm{E} \rangle = k_{\mathrm{INI},a} \bar{g}_a \langle n_a \mid \mathrm{E} \rangle, \quad \langle m_b \mid \mathrm{E} \rangle = k_{\mathrm{INI},b} \bar{g}_b \langle n_b \mid \mathrm{E} \rangle, \tag{S54}$$

$$\sigma^2_{m_a|\mathrm{E}} = k^2_{\mathrm{INI},a} T^2_{\mathrm{RES}} \bar{g}^2_a \sigma^2_{n_a|\mathrm{E}} + k_{\mathrm{INI},a} T_{\mathrm{RES}} \overline{g^2_a} \langle n_a \mid \mathrm{E} \rangle, \quad \sigma^2_{m_b|\mathrm{E}} = k^2_{\mathrm{INI},b} T^2_{\mathrm{RES}} \bar{g}^2_b \sigma^2_{n_b|\mathrm{E}} + k_{\mathrm{INI},b} T_{\mathrm{RES}} \overline{g^2_b} \langle n_b \mid \mathrm{E} \rangle, \tag{S55}$$

$$\mathrm{Cov}(m_a, m_b \mid \mathrm{E}) = \bar{g}_a \bar{g}_b k_{\mathrm{INI},a} k_{\mathrm{INI},b} f_\beta(\langle n_a \mid \mathrm{E} \rangle, \gamma). \tag{S56}$$

Meanwhile, the covariance caused by the extrinsic fluctuation is given by:

$$\mathrm{Cov}(\langle m_a \mid \mathrm{E} \rangle, \langle m_b \mid \mathrm{E} \rangle) = \bar{g}_a \bar{g}_b k_{\mathrm{INI},a} k_{\mathrm{INI},b} \mathrm{Cov}(\langle n_a \mid \mathrm{E} \rangle, \langle n_b \mid \mathrm{E} \rangle) = \bar{g}_a \bar{g}_b k_{\mathrm{INI},a} k_{\mathrm{INI},b} \sigma^2_{\langle n_a|\mathrm{E}\rangle} \left. \frac{\partial h_\beta}{\partial n} \right|_{\langle n_a \rangle}. \tag{S57}$$

Combining **Eqs. (S56)-(S57)**, we express total covariance as:

$$\mathrm{Cov}(m_a, m_b) \approx \bar{g}_a \bar{g}_b k_{\mathrm{INI},a} k_{\mathrm{INI},b} \left[ f_\beta(\langle n_a \rangle, \gamma) + \left( \frac{1}{2} \left. \frac{\partial^2 f_\beta}{\partial n^2} \right|_{\langle n_a \rangle} + \left. \frac{\partial h_\beta}{\partial n} \right|_{\langle n_a \rangle} \right) \sigma^2_{\langle n_a|\mathrm{E}\rangle} \right]. \tag{S58}$$

Therefore, we can determine $\gamma$ from the partial slope of $\mathrm{Cov}(m_a, m_b)$ with respect to $\sigma^2_{\langle m_a|\mathrm{E}\rangle}$, i.e.,

$$\left. \frac{\partial \mathrm{Cov}(m_a, m_b)}{\partial \sigma^2_{\langle m_a|\mathrm{E}\rangle}} \right|_{\langle n_a \rangle} \approx \frac{\bar{g}_b k_{\mathrm{INI},b}}{\bar{g}_a k_{\mathrm{INI},a}} \left( \frac{1}{2} \left. \frac{\partial^2 f_\beta}{\partial n^2} \right|_{\langle n_a \rangle} + \left. \frac{\partial h_\beta}{\partial n} \right|_{\langle n_a \rangle} \right), \tag{S59}$$

where $\langle n_a \rangle$, $\bar{g}_a k_{\mathrm{INI},a}$, and $\bar{g}_b k_{\mathrm{INI},b}$ are estimated from the total noise of $m_a$ and $m_b$.

**Supplemental Material**

When applying the above method to analyze nascent RNA signals from alternative *hb* promoters, we can estimate $\beta$ from the ratio between $\langle n_b \mid x \rangle$ and $\langle n_a \mid x \rangle$ measured at the low-expression region of the embryo, i.e.,

$$\lim_{n_a \to 0} \frac{\langle n_b \mid x \rangle}{\langle n_a \mid x \rangle} = \lim_{\frac{k_{\mathrm{ON},a}}{k_{\mathrm{OFF},a}} \to 0} \frac{\beta + \beta\gamma \left( \frac{k_{\mathrm{ON},a}}{k_{\mathrm{OFF},a}} \right)^2}{1 + \beta\gamma \left( \frac{k_{\mathrm{ON},a}}{k_{\mathrm{OFF},a}} \right)^2} = \beta. \quad \text{(S60)}$$

Our experimental data show that such ratio for *hb* P1 and P2 stays around 0.074 in the posterior part of the embryo (>0.5 EL), where the *hb* expression is close to zero (**Fig. S3(c)**), suggesting that $\beta \approx 0.074$.

#### 13. Generalization of the framework to diverse coupling mechanisms

Although our method was derived from a simple two-state transcription model with a specific $k_{\mathrm{ON}}$-coupling kinetics, most of the results are generally applicable to other models with diverse coupling mechanisms. Below, we assess its applicability to several biologically relevant transcriptional coupling mechanisms through theoretical analysis and stochastic simulations.

##### 13.1. TF-sharing mechanism

A common form of gene-gene coupling arises from competition for shared transcription factors (TFs)[33,34]. To model this mechanism, we consider a simple telegraph model of two identical genes ($i$ = $a$, $b$), each requiring the binding of a common TF species for activation (**Fig. S4(a)**). Specifically, we assume that the total number of TF molecules in the cell is a finite number, denoted by $N$. Each TF molecule can either stay in the cytoplasm (of volume $V$) or bind to a specific regulatory site on either gene to activate transcription. The activation rate ($k_{\mathrm{ON}}$) of a gene is proportional to the concentration of unbound TFs in the cytoplasm. Thus, when both genes are OFF, $k_{\mathrm{ON}} = k_0 N / V$, where $k_0$ is a proportionality constant. In contrast, when one gene turns ON, it changes the activation rate of the other gene to $k_{\mathrm{ON}} = k_0 (N-1) / V$.

Clearly, this model represents a special case of our original $k_{\mathrm{ON}}$-coupling model with $\gamma = 1 - 1/N$. Therefore, all main-text results remain valid. Note that the above derivation assumes a simple case with a single TF binding site on the regulatory sequence of the gene. In case multiple TF binding sites are present, the expression for $\gamma(N)$ will be different.

To verify this, we performed Gillespie simulations of this TF-sharing model under three TF abundance conditions: $N = 1$ (extremely scarce), $N = 2$ (moderate), and $N = 10^3$ (extremely abundant) (**Fig. S4(b)**). Other kinetic parameters were set as follows: $V = 1$, $k_0 = 0.01$–$0.05$ min$^{-1}$, $k_{\mathrm{OFF}} = 0.05$ min$^{-1}$, $k_{\mathrm{INI}} = 50$ min$^{-1}$, $T_{\mathrm{RES}} = 2$ min, and $g(\tau) = -\tau$. As expected, the resulting relationships among $C_{m_a,m_b}$, $\langle n \rangle$, and $\sigma^2_{\langle n|\mathrm{E}\rangle}$ aligned with the cases of $\gamma = 0$, $0 < \gamma < 1$, and $\gamma = 1$ in **Fig. 3(a)**, respectively. Moreover, applying our noise decomposition framework to the simulation data with $N = 2$ yielded an estimated $\gamma = 0.47$, in excellent agreement with the theoretical value $\gamma = 0.5$ (**Fig. S4(c)**), validating the effectiveness of our method.

##### 13.2. $k_{\mathrm{ON}}$-coupling in prokaryotes with co-transcriptional degradation

In prokaryotes, mRNAs often undergo rapid degradation due to the absence of a nuclear membrane and the tight coupling between transcription and translation. In particular, degradation can begin during transcription elongation, acting on nascent transcripts before completion[28,29]. To examine how this process influences correlated noise between genes, we extend our original $k_{\mathrm{ON}}$-coupling model to incorporate co-transcriptional degradation process (**Fig. S4(d)**). Specifically, we assume that each nascent RNA molecule, once initiated, can randomly start degradation with a rate $k_{\mathrm{d}}$. Once the degradation starts, it proceeds from the 5' to the 3' end of the RNA at the same speed as the transcription elongation speed $V_{\mathrm{EL}}$[28,29]. The release of nascent RNA from the gene is unaffected by degradation.

Although analytically solving this model is difficult, it is evident that the gene-state dynamics remain unchanged from the original $k_{\mathrm{ON}}$-coupling model, i.e., **Eqs. (2)-(4)** in the main text remain valid. Moreover, in the limit of slow gene-state transitions ($k_{\mathrm{ON}}$ & $k_{\mathrm{OFF}} \to 0$), the linear relationship between nascent RNA statistics and gene-state statistics should still hold, albeit with modified proportionality constants. Therefore, our noise decomposition framework, including **Eq. (12)** and the estimation of $\langle n \rangle$ from $\eta_m^2$, remains applicable.

To verify these theoretical arguments, we conducted Gillespie simulations for three different coupling strengths: $\gamma = 0.5$, 1, and 2 (**Fig. S4(e)**). Other kinetic parameters were set as follows: $k_{\mathrm{ON}} = 0.01$–$0.05$ min$^{-1}$, $k_{\mathrm{OFF}} = 0.05$ min$^{-1}$, $k_{\mathrm{INI}} = 50$ min$^{-1}$, $T_{\mathrm{RES}} = 2$ min, $g(\tau) = -\tau$, and $k_{\mathrm{d}} = 0.5$ min$^{-1}$. As expected, the relationship among $C_{m_a,m_b}$, $\langle n \rangle$, and $\sigma^2_{\langle n|\mathrm{E}\rangle}$ for every $\gamma$ value aligned with the corresponding cases in **Fig. 3(a)**. Moreover, applying our noise decomposition framework accurately recovered the corresponding $\gamma$ values (**Fig. S4(f)**). These results confirm the applicability of our noise decomposition framework to genes with co-transcriptional degradation.

##### 13.3. $k_{\mathrm{OFF}}$-coupling mechanism

Complementary to $k_{\mathrm{ON}}$-coupling, our model can be modified to describe a coupling mechanism mediated through gene inactivation (**Fig. S4(g)**). Specifically, we consider that, if one gene copy turns OFF, it modulates the deactivation rate of its sister copy by a factor $\gamma$, i.e., $k_{\mathrm{OFF}} \to \gamma k_{\mathrm{OFF}}$. In this case, the gene-state transition rate matrix becomes

$$\mathbf{K} = \begin{bmatrix} -2k_{\mathrm{ON}} & \gamma k_{\mathrm{OFF}} & \gamma k_{\mathrm{OFF}} & 0 \\ k_{\mathrm{ON}} & -k_{\mathrm{ON}} - \gamma k_{\mathrm{OFF}} & 0 & k_{\mathrm{OFF}} \\ k_{\mathrm{ON}} & 0 & -k_{\mathrm{ON}} - \gamma k_{\mathrm{OFF}} & k_{\mathrm{OFF}} \\ 0 & k_{\mathrm{ON}} & k_{\mathrm{ON}} & -2k_{\mathrm{OFF}} \end{bmatrix}. \tag{S61}$$

Under fixed environmental conditions, it leads to a steady-state distribution of gene states $n_a$ and $n_b$:

$$\tilde{\mathbf{P}}_0 = \frac{1}{\gamma\left(k_{\text{OFF}}/k_{\text{ON}}\right)^2 + 2\left(k_{\text{OFF}}/k_{\text{ON}}\right)+1}\begin{bmatrix}\gamma\left(k_{\text{OFF}}/k_{\text{ON}}\right)^2 \\ k_{\text{OFF}}/k_{\text{ON}} \\ k_{\text{OFF}}/k_{\text{ON}} \\ 1\end{bmatrix}. \tag{S62}$$

Notably, by exchanging the ON and OFF states (along with the corresponding $k_{\text{ON}}$ and $k_{\text{OFF}}$), this expression reduces to **Eq. (2)** in the main text. Since $C_{n_a,n_b|\text{E}}$ is invariant under such ON-OFF transformation, all results derived for the $k_{\text{ON}}$-coupling model, including the expression of $f$ (**Eq. (4)**), remain valid. In essence, $k_{\text{ON}}$- and $k_{\text{OFF}}$-coupling models are equivalent in their ability to generate correlated noise.

To verify these theoretical arguments, we performed Gillespie simulations under three different coupling strengths: $\gamma = 0.5$, 1, and 2 (**Fig. S4(h)**). Other kinetic parameters were set as follows: $k_{\text{ON}} = 0.01–0.05$ min$^{-1}$, $k_{\text{OFF}} = 0.05$ min$^{-1}$, $k_{\text{INI}} = 50$ min$^{-1}$, $T_{\text{RES}} = 2$ min, and $g(\tau) = -\tau$. As expected, the relationship among $C_{m_a,m_b}$, $\langle n \rangle$, and $\sigma^2_{\langle n|\text{E}\rangle}$ for every $\gamma$ value matched the corresponding cases in **Fig. 3(a)**. Furthermore, applying our noise decomposition framework accurately recovered the corresponding $\gamma$ value (**Fig. S4(i)**), validating the effectiveness of our method.

##### 13.4. Coupled multi-state models

Beyond the simple two-state model, transcription kinetics is sometimes described using more complex multi-state models that account for intermediate steps in transcriptional regulation, such as chromatin remodeling[35,36], pre-initiation complex (PIC) formation[37,38], and promoter conformation changes[39]. In these models, a gene may fluctuate among multiple active (ON) and/or inactive (OFF) transcription states, with distinct transition rates[7,40,41]. To evaluate the applicability of our method in such scenarios, we extend the original two-state model to a three-state version and examine coupling effects in two representative cases:

**Supplemental Material**

**13.4.1. Coupled "OFF-OFF-ON" model**

A typical type of the three-state model comprises two OFF states (States 0 and 1) and one ON state (State 2), with sequential stochastic transitions between adjacent states at rates $k_{ij}$ ($i$, $j$ = 0, 1, 2 and $|i-j| = 1$) (**Fig. S4(j)**). In this framework, the gene must transition from a basal OFF state to an intermediate OFF state before reaching the ON state. This model captures the experimentally observed non-exponential distribution of inactive periods[40–42], which may reflect essential intermediate steps in gene activation, such as chromatin remodeling or PIC formation[35–38]. To introduce gene-gene coupling into this model, we assume that, if one gene copy is in the ON state, it modulates the transition rate from the intermediate OFF state (State 1) to the ON state (State 2) of its sister copy by a factor $\gamma$, i.e., $k_{12} \to \gamma k_{12}$ (**Fig. S4(j)**). The master equation of this model remains **Eq. (A1)**, with all matrix components expanded to accommodate the higher-dimensional state space. Specifically,

$$\mathbf{P}(m_a, m_b) = \begin{bmatrix} P(n_a=0, n_b=0, m_a, m_b) \\ P(n_a=1, n_b=0, m_a, m_b) \\ P(n_a=2, n_b=0, m_a, m_b) \\ P(n_a=0, n_b=1, m_a, m_b) \\ P(n_a=1, n_b=1, m_a, m_b) \\ P(n_a=2, n_b=1, m_a, m_b) \\ P(n_a=0, n_b=2, m_a, m_b) \\ P(n_a=1, n_b=2, m_a, m_b) \\ P(n_a=2, n_b=2, m_a, m_b) \end{bmatrix},$$

$$\mathbf{K} = \begin{bmatrix} -2k_{01} & k_{10} & 0 & k_{10} & 0 & 0 & 0 & 0 & 0 \\ k_{01} & -k_{10}-k_{01}-k_{12} & k_{21} & 0 & k_{10} & 0 & 0 & 0 & 0 \\ 0 & k_{12} & -k_{21}-k_{01} & 0 & 0 & k_{10} & 0 & 0 & 0 \\ k_{01} & 0 & 0 & -k_{10}-k_{01}-k_{12} & k_{10} & 0 & k_{21} & 0 & 0 \\ 0 & k_{01} & 0 & k_{01} & -2k_{10}-2k_{12} & k_{21} & 0 & k_{21} & 0 \\ 0 & 0 & k_{01} & 0 & k_{12} & -k_{10}-k_{21}-\gamma k_{12} & 0 & 0 & k_{21} \\ 0 & 0 & 0 & k_{12} & 0 & 0 & -k_{21}-k_{01} & k_{10} & 0 \\ 0 & 0 & 0 & 0 & k_{12} & 0 & k_{01} & -k_{10}-k_{21}-\gamma k_{12} & k_{21} \\ 0 & 0 & 0 & 0 & 0 & \gamma k_{12} & 0 & \gamma k_{12} & -2k_{21} \end{bmatrix},$$

$$\mathbf{K}_{\mathrm{INI},a}=\begin{bmatrix}0&0&0&0&0&0&0&0&0\\0&0&0&0&0&0&0&0&0\\0&0&k_{\mathrm{INI}}&0&0&0&0&0&0\\0&0&0&0&0&0&0&0&0\\0&0&0&0&0&0&0&0&0\\0&0&0&0&0&k_{\mathrm{INI}}&0&0&0\\0&0&0&0&0&0&0&0&0\\0&0&0&0&0&0&0&0&0\\0&0&0&0&0&0&0&0&k_{\mathrm{INI}}\end{bmatrix},\text{ and }\mathbf{K}_{\mathrm{INI},b}=\begin{bmatrix}0&0&0&0&0&0&0&0&0\\0&0&0&0&0&0&0&0&0\\0&0&0&0&0&0&0&0&0\\0&0&0&0&0&0&0&0&0\\0&0&0&0&0&0&0&0&0\\0&0&0&0&0&0&0&0&0\\0&0&0&0&0&0&k_{\mathrm{INI}}&0&0\\0&0&0&0&0&0&0&k_{\mathrm{INI}}&0\\0&0&0&0&0&0&0&0&k_{\mathrm{INI}}\end{bmatrix}.$$

Under this formulation, the general expressions for the mean, variance, and covariance of nascent RNA (**Eqs. (5)-(7)**, **(13)**, **(A2)-(A4)**) stay structurally unchanged. Although the explicit form of $\langle n\,|\,\mathrm{E}\rangle$ as a function of $k_{ij}$ differs from the two-state model, a computer-assisted analytical derivation (MATLAB) reveals that the relationship between $C_{n_a,n_b|\mathrm{E}}$, $\langle n\,|\,\mathrm{E}\rangle$ and $\gamma$, i.e., $f(n,\gamma)$, is consistent with **Eq. (4)**. Therefore, our noise decomposition framework remains applicable.

To verify these theoretical arguments, we conducted Gillespie simulations for three different coupling strengths: $\gamma = 0.5$, 1, and 2 (**Fig. S4(k)**). The other kinetic parameters were set as follows: $k_{01} = 0.01$–$0.08$ min$^{-1}$, $k_{12} = 0.01$–$0.08$ min$^{-1}$, $k_{10} = 0.05$ min$^{-1}$, $k_{21} = 0.05$ min$^{-1}$, $k_{\mathrm{INI}} = 50$ min$^{-1}$, $T_{\mathrm{RES}} = 2$ min, and $g(\tau) = -\tau$. As expected, the relationship among $C_{m_a,m_b}$, $\langle n\rangle$, and $\sigma^2_{\langle n|\mathrm{E}\rangle}$ for every $\gamma$ value matched the corresponding cases in **Fig. 3(a)**. Furthermore, applying our noise decomposition framework accurately recovered the corresponding $\gamma$ values (**Fig. S4(l)**), validating the effectiveness of our method for the coupled "OFF-OFF-ON" gene pairs.

**Supplemental Material**

**13.4.2. Coupled "OFF-ON-ON" model**

Another common form of the three-state model comprises one OFF state (State 0) and two ON states (States 1 and 2), with sequential stochastic transitions between adjacent states at rates $k_{ij}$ ($i, j = 0, 1, 2$ and $|i-j| = 1$) (**Fig. S4(m)**). This "OFF-ON-ON" model captures distinct conformations of the active promoter, which may be essential in certain biological systems[7,39]. To introduce gene-gene coupling into this model, we assume that, if one gene copy is in either of the ON states, it modulates the transition rate from the OFF state (State 0) to the intermediate ON state (State 1) of its sister copy by a factor $\gamma$, i.e., $k_{01} \rightarrow \gamma k_{01}$ (**Fig. S4(m)**). For simplicity, we assume that both ON states share the same transcription initiation rate, i.e., $k_{\mathrm{INI},1} = k_{\mathrm{INI},2}$. The master equation of this "OFF-ON-ON" model remains **Eq. (A1)**, with

$$\mathbf{K} = \begin{bmatrix} -2k_{01} & k_{10} & 0 & k_{10} & 0 & 0 & 0 & 0 & 0 \\ k_{01} & -\gamma k_{01} - k_{10} - k_{12} & k_{21} & 0 & k_{10} & 0 & 0 & 0 & 0 \\ 0 & k_{12} & -\gamma k_{01} - k_{21} & 0 & 0 & k_{10} & 0 & 0 & 0 \\ k_{01} & 0 & 0 & -\gamma k_{01} - k_{10} - k_{12} & k_{10} & 0 & k_{21} & 0 & 0 \\ 0 & \gamma k_{01} & 0 & \gamma k_{01} & -2k_{10} - 2k_{12} & k_{21} & 0 & k_{21} & 0 \\ 0 & 0 & \gamma k_{01} & 0 & k_{12} & -k_{10} - k_{21} - k_{12} & 0 & 0 & k_{21} \\ 0 & 0 & 0 & k_{12} & 0 & 0 & -\gamma k_{01} - k_{21} & k_{10} & 0 \\ 0 & 0 & 0 & 0 & k_{12} & 0 & \gamma k_{01} & -k_{10} - k_{21} - k_{12} & k_{21} \\ 0 & 0 & 0 & 0 & 0 & k_{12} & 0 & k_{12} & -2k_{21} \end{bmatrix},$$

$$\mathbf{K}_{\mathrm{INI},a} = \begin{bmatrix} 0 & 0 & 0 & 0 & 0 & 0 & 0 & 0 & 0 \\ 0 & k_{\mathrm{INI}} & 0 & 0 & 0 & 0 & 0 & 0 & 0 \\ 0 & 0 & k_{\mathrm{INI}} & 0 & 0 & 0 & 0 & 0 & 0 \\ 0 & 0 & 0 & 0 & 0 & 0 & 0 & 0 & 0 \\ 0 & 0 & 0 & 0 & k_{\mathrm{INI}} & 0 & 0 & 0 & 0 \\ 0 & 0 & 0 & 0 & 0 & k_{\mathrm{INI}} & 0 & 0 & 0 \\ 0 & 0 & 0 & 0 & 0 & 0 & 0 & 0 & 0 \\ 0 & 0 & 0 & 0 & 0 & 0 & 0 & k_{\mathrm{INI}} & 0 \\ 0 & 0 & 0 & 0 & 0 & 0 & 0 & 0 & k_{\mathrm{INI}} \end{bmatrix}, \text{ and } \mathbf{K}_{\mathrm{INI},b} = \begin{bmatrix} 0 & 0 & 0 & 0 & 0 & 0 & 0 & 0 & 0 \\ 0 & 0 & 0 & 0 & 0 & 0 & 0 & 0 & 0 \\ 0 & 0 & 0 & 0 & 0 & 0 & 0 & 0 & 0 \\ 0 & 0 & 0 & k_{\mathrm{INI}} & 0 & 0 & 0 & 0 & 0 \\ 0 & 0 & 0 & 0 & k_{\mathrm{INI}} & 0 & 0 & 0 & 0 \\ 0 & 0 & 0 & 0 & 0 & k_{\mathrm{INI}} & 0 & 0 & 0 \\ 0 & 0 & 0 & 0 & 0 & 0 & k_{\mathrm{INI}} & 0 & 0 \\ 0 & 0 & 0 & 0 & 0 & 0 & 0 & k_{\mathrm{INI}} & 0 \\ 0 & 0 & 0 & 0 & 0 & 0 & 0 & 0 & k_{\mathrm{INI}} \end{bmatrix}.$$

Under this formulation, the general expressions for the mean, variance, and covariance of nascent RNA (**Eqs. (5)-(7)**, **(13)**, **(A2)-(A4)**), as well as the expression of $f$ (**Eq. (4)**), stay unchanged (confirmed through a computer-assisted analytical derivation (MATLAB)). Therefore, our noise decomposition framework remains applicable to the coupled "OFF-ON-ON" model.

To verify these theoretical arguments, we conducted Gillespie simulations for three different coupling strengths: $\gamma = 0.5$, 1, and 2 (**Fig. S4(n)**). The other kinetic parameters were set as follows: $k_{01} = 0.01$–$0.05$ min$^{-1}$, $k_{12} = 0.01$–$0.05$ min$^{-1}$, $k_{10} = 0.05$ min$^{-1}$, $k_{21} = 0.05$ min$^{-1}$, $k_{\mathrm{INI}} = 50$ min$^{-1}$, $T_{\mathrm{RES}} = 2$ min, and $g(\tau) = -\tau$. As expected, the relationship among $C_{m_a,m_b}$, $\langle n \rangle$, and $\sigma^2_{\langle n|\mathrm{E}\rangle}$ for every $\gamma$ value matched the corresponding cases in **Fig. 3(a)**. Furthermore, applying our noise decomposition framework accurately recovered the corresponding $\gamma$ values (**Fig. S4(o)**), validating the effectiveness of our method for the coupled "OFF-ON-ON" gene pairs.

##### 13.5. RNAP-sharing mechanism

Besides coupling at the gene-state transition level, a prevalent form of gene-gene coupling arises at the transcription initiation stage. Biologically, this can occur through a competition between genes for a limited pool of RNA polymerases (RNAPs)[43] (**Fig. S5(a)**). Specifically, we assume that two genes ($i = a$, $b$) independently switch between ON and OFF states, while sharing a finite number ($N_{\mathrm{P0}}$) of RNAP molecules within the cell. Each RNAP molecule can either stay in the cytoplasm (of volume $V$) or engage with a gene to initiate transcription. The transcription initiation rate ($k_{\mathrm{INI}}$) of a gene is proportional to the cytoplasmic RNAP concentration, i.e.:

$$k_{\mathrm{INI}} = k_{\mathrm{INI0}}(N_{\mathrm{P0}} - N_{\mathrm{P},a} - N_{\mathrm{P},b}) / V, \tag{S63}$$

where $N_{\mathrm{P},i}$ is the number of RNAPs currently engaged with gene $i$, and $k_{\mathrm{INI0}}$ is a proportionality constant. Note that, in this system, the two genes are independent at the gene-state level, i.e., $C_{n_a,n_b|\mathrm{E}} = 0$, while their $C_{m_a,m_b|\mathrm{E}}$ arises from the fluctuations in $k_{\mathrm{INI}}$ over time.

Since $k_{\mathrm{INI}}$ depends on $N_{\mathrm{P},i}$ (and therefore, on $m_i$), such system is analytically intractable. Nevertheless, the intrinsic nascent RNA covariance can be generally written as $C_{m_a,m_b|\mathrm{E}} = \bar{g}^2 k_{\mathrm{INI0}}^2 f(\langle n\,|\,\mathrm{E}\rangle, N_{\mathrm{P0}})$, where $N_{\mathrm{P0}}$ is a measure of the coupling strength, though the expression of $f$ may differ from **Eq. (4)** in the main text.

To gain further analytical insight into this mechanism, we simplify the model into a more tractable form. I.e., instead of modeling the dependence of $k_{\mathrm{INI}}$ on the actual number of RNAP molecules, we assume that the transcription initiation rate of each gene copy depends solely on the activation status of its sister copy. Specifically, the baseline initiation rate ($k_{\mathrm{INI}}$) is constant for both gene copies. If one gene copy turns ON, it modulates the initiation rate of its sister copy by a factor $\gamma$, i.e., $k_{\mathrm{INI}} \to \gamma k_{\mathrm{INI}}$ (**Fig. S5(b)**). Since the two gene copies compete for RNAP, $\gamma$ is typically less than one. The master equation of this model remains **Eq. (A1)**, with

$$\mathbf{K} = \begin{bmatrix} -2k_{\mathrm{ON}} & k_{\mathrm{OFF}} & k_{\mathrm{OFF}} & 0 \\ k_{\mathrm{ON}} & -k_{\mathrm{ON}} - k_{\mathrm{OFF}} & 0 & k_{\mathrm{OFF}} \\ k_{\mathrm{ON}} & 0 & -k_{\mathrm{ON}} - k_{\mathrm{OFF}} & k_{\mathrm{OFF}} \\ 0 & k_{\mathrm{ON}} & k_{\mathrm{ON}} & -2k_{\mathrm{OFF}} \end{bmatrix}, \quad \mathbf{K}_{\mathrm{INI},a} = \begin{bmatrix} 0 & 0 & 0 & 0 \\ 0 & k_{\mathrm{INI}} & 0 & 0 \\ 0 & 0 & 0 & 0 \\ 0 & 0 & 0 & \gamma k_{\mathrm{INI}} \end{bmatrix}, \text{ and}$$

$$\mathbf{K}_{\mathrm{INI},b} = \begin{bmatrix} 0 & 0 & 0 & 0 \\ 0 & 0 & 0 & 0 \\ 0 & 0 & k_{\mathrm{INI}} & 0 \\ 0 & 0 & 0 & \gamma k_{\mathrm{INI}} \end{bmatrix}.$$

Following **Appendix A**, we find that,

$$\langle m\,|\,\mathrm{E}\rangle = \bar{g} k_{\mathrm{INI}} \left[ n + (\gamma - 1) n^2 \right], \tag{S64}$$

$$\sigma_{m|\mathrm{E}}^2 = \bar{g}^2 k_{\mathrm{INI}}^2 \left[ n(1-n) + (\gamma^2 - 1) n^2 - 2(\gamma - 1) n^3 - (\gamma - 1)^2 n^4 \right] + \overline{g^2} k_{\mathrm{INI}} \left[ n(1-n) + \gamma n^2 \right], \tag{S65}$$

where $n \equiv \langle n\,|\,\mathrm{E}\rangle$. Here, unlike **Eqs. (5)** and **(7)** in the main text, both the mean and variance of nascent RNA depend on $\gamma$ and exhibit complex relationships with $\langle n\,|\,\mathrm{E}\rangle$ and $\sigma_{n|\mathrm{E}}^2$. However, for small $n$, by neglecting higher-order terms, we may still apply **Eq. (13)** to estimate $\langle n \mid \mathrm{E} \rangle$ from $\eta_m^2$. Moreover, the intrinsic nascent RNA covariance can be written as $C_{m_a,m_b|\mathrm{E}} = \bar{g}^2 k_{\mathrm{INI}}^2 f(\langle n \mid \mathrm{E} \rangle, \gamma)$, where

$$f(n,\gamma) = n^2(1-n)(\gamma-1)(\gamma-n+n\gamma+1). \tag{S66}$$

This expression is different from **Eq. (4)** in the main text, highlighting that the form of $f$ depends on the underlying coupling mechanism, and can thus be used to infer the mechanism from experimental data.

To verify the above results, we conducted Gillespie simulations for this RNAP-sharing model under varying levels of RNAP abundance: $N_{\mathrm{P0}}$ = 1 (extremely scarce), $N_{\mathrm{P0}}$ = 10 (moderate), and $N_{\mathrm{P0}}$ = $10^4$ (extremely abundant) (**Fig. S5(c)**). Other kinetic parameters were set as follows: $V$ = 1, $k_{\mathrm{ON}}$ = 0.01–0.05 min$^{-1}$, $k_{\mathrm{OFF}}$ = 0.05 min$^{-1}$, $k_{\mathrm{INI0}}$ = 50 min$^{-1}$, $T_{\mathrm{RES}}$ = 2 min, and $g(\tau) = -\tau$. The resulting relationships between $C_{m_a,m_b}$, $\langle n \rangle$, and $\sigma^2_{\langle n|\mathrm{E}\rangle}$ qualitatively matched the cases of $\gamma = 0$, $0 < \gamma < 1$, and $\gamma = 1$ in **Fig. 3(a)**, validating the general idea of our model. However, directly applying **Eq. (4)** from the main text failed to yield a common solution for $\gamma$ across different $\langle n \rangle$ values (**Fig. S5(d)**). In contrast, applying **Eq. (S66)** successfully identified an equivalent $\gamma$ for the simplified model (**Fig. S5(e)**). These results demonstrate the ability of our method to distinguish between different coupling mechanisms. In case the precise coupling mechanism is unknown, this capability enables the exploration of plausible coupling scenarios by testing various candidate forms of $f(n,\gamma)$.

##### 13.6. Feedback mechanisms

Beyond direct interactions, gene-gene coupling can also arise from feedback regulation mediated by the genes' own RNA or protein products[44,45]. Since the accumulation of these products typically occurs over a time scale of hours to days, their regulatory effects are effectively constant within the much shorter time window of nascent RNA transcription. Consequently, feedback-induced covariance is generally negligible at the nascent RNA level.

Nevertheless, to test the applicability of our method to such mechanisms, we consider a scenario where feedback is directly mediated by the genes' mRNA products. In this case, the coupling effect may become detectable at the level of total (rather than nascent) cellular mRNA. Specifically, we examine the following two types of regulation:

##### 13.6.1. Autoregulation

A simple form of feedback regulation that can couple two identical genes is autoregulation, in which the mRNA product from either gene can activate or repress the transcription of both genes (**Fig. S5(f)**). To model this mechanism, we start with a standard two-state telegraph model for cellular mRNA levels produced by a pair of identical genes. Compared to the nascent RNA model, this model overlooks the RNA production process (i.e., synthesis, post-synthesis residence, and release), but includes a constant-rate ($k_D$) degradation of each mRNA molecule. We assume that the activation rate of each gene is modulated by the total mRNA level from both genes, following a Hill function:

$$k_{ON} = \left(k_{ON1} - k_{ON0}\right)\frac{(m_a + m_b)^h}{(m_a + m_b)^h + m_0^h} + k_{ON0}. \tag{S67}$$

Here, $k_{ON0}$ and $k_{ON1}$ represent the basal and maximum activation rates, respectively; $m_0$ is the characteristic mRNA level corresponding to half-maximal regulatory effect; and $h$ is the Hill coefficient reflecting the cooperativity of autoregulation. $k_{ON1}$, $m_0$, and $h$ all influence the strength of coupling between the two genes. In particular, $h > 0$ corresponds to auto-activation, while and $h < 0$ corresponds to auto-repression.

The master equation of this system is written as:

$$\frac{d\mathbf{P}(m_a,m_b)}{dt} = (\mathbf{K}-\mathbf{K}_{\mathrm{INI},a}-\mathbf{K}_{\mathrm{INI},b}-(m_a+m_b)k_{\mathrm{D}})\mathbf{P}(m_a,m_b)$$
$$+\mathbf{K}_{\mathrm{INI},a}\mathbf{P}(m_a-1,m_b)+\mathbf{K}_{\mathrm{INI},b}\mathbf{P}(m_a,m_b-1)$$
$$+(m_a+1)k_{\mathrm{D}}\mathbf{P}(m_a+1,m_b)+(m_b+1)k_{\mathrm{D}}\mathbf{P}(m_a,m_b+1). \quad \text{(S68)}$$

Note that, without modeling nascent RNA production process, this equation directly describes the experimentally observable $m_a$, $m_b$, and $t$, instead of $\mathcal{m}_a$, $\mathcal{m}_b$, and $\tau$. Similar to the RNAP-sharing mechanism, the dependence of $k_{\mathrm{ON}}$ on $m_i$ makes **Eq. (S68)** analytically intractable. Nevertheless, for given $k_{\mathrm{ON1}}$ and $m_0$, the intrinsic covariance of cellular mRNA can, in principle, be expressed as $C_{m_a,m_b|\mathrm{E}} = \bar{g}^2 k_{\mathrm{INI}}^2 f(\langle n\,|\,\mathrm{E}\rangle, h)$, although the exact form of $f$ differs from that in **Eq. (4)** of the main text. Therefore, the overall logic of our noise decomposition framework, including **Eqs. (12)** and **(13)**, should remain valid.

To test this, we performed stochastic simulations for this autoregulation model under various Hill coefficients: $h$ = 0, and ±1(**Fig. S5(g)**). Other kinetic parameters were set as follows: $k_{\mathrm{ON0}} = 0.01\ \mathrm{min}^{-1}$, $k_{\mathrm{ON1}} = 0.01\text{–}0.05\ \mathrm{min}^{-1}$, $m_0 = 5$, $k_{\mathrm{OFF}} = 0.05\ \mathrm{min}^{-1}$, $k_{\mathrm{INI}} = 50\ \mathrm{min}^{-1}$, $k_{\mathrm{D}} = 0.5\ \mathrm{min}^{-1}$, $T_{\mathrm{RES}} = 2$ min, and $g(\tau) = -\tau$. The resulting relationships between $C_{m_a,m_b}$, $\langle n\rangle$, and $\sigma^2_{\langle n|\mathrm{E}\rangle}$ qualitatively matched **Fig. 3(a)**, i.e., $h = -1$ (negative feedback) correspond to $0 < \gamma < 1$ (negative coupling), $h = 0$ (no feedback) corresponds to $\gamma = 1$ (no coupling), and $h = 1$ (positive feedback) correspond to $\gamma > 1$ (positive coupling). These results support the overall idea of our method and reveal a potential for quantifying autoregulation-induced coupling, provided that an appropriate $f(n,\gamma)$ is used.

##### 13.6.2. Toggle switch

In addition to autoregulation, feedback regulation can also occur through a toggle switch mechanism, where the mRNA product from each gene represses the transcription of the other gene (**Fig. S5(h)**). To model this mechanism, we consider a two-state telegraph model for cellular mRNA levels of two distinct genes. The activation rate of each gene is regulated by the total mRNA level of its counterpart, following a Hill function:

$$k_{\mathrm{ON},i} = \left(k_{\mathrm{ON1}} - k_{\mathrm{ON0}}\right)\frac{m_0^h}{m_j^h + m_0^h} + k_{\mathrm{ON0}}. \tag{S69}$$

where $i \neq j$, and other parameters are defined as in **Eq. (S67)**.

The master equation of this system remains **Eq. (S68)**, which is analytically intractable. However, with appropriate $f(\langle n \mid \mathrm{E}\rangle, h)$, the core principle of our noise decomposition method remains applicable. To verify this, we conducted stochastic simulations under varying values of Hill coefficient: $h$ = 0, 1, and 5 (**Fig. S5(i)**). Other kinetic parameters were set as follows: $k_{\mathrm{ON0}}$ = 0.01 min$^{-1}$, $k_{\mathrm{ON1}}$ = 0.01–0.05 min$^{-1}$, $m_0$ = 5, $k_{\mathrm{OFF}}$ = 0.05 min$^{-1}$, $k_{\mathrm{INI}}$ = 50 min$^{-1}$, $k_{\mathrm{D}}$ = 1 min$^{-1}$, $T_{\mathrm{RES}}$ = 2 min, and $g(\tau) = -\tau$. The resulting relationships between $C_{m_a,m_b}$, $\langle n \rangle$, and $\sigma^2_{\langle n|\mathrm{E}\rangle}$ qualitatively matched **Fig. 3(a)**, with $h$ = 0, 1, and 5 corresponding to no coupling ($\gamma$ = 1), weak negative coupling ($0 < \gamma < 1$), and strong negative coupling ($\gamma \to 0$), respectively. These results support the overall idea of our method and reveal a potential for quantifying toggle-switch-induced coupling, provided that an appropriate $f(n,g)$ is used.

#### Supplemental Material

**Supplemental Material**

28. Chen, H., Shiroguchi, K., Ge, H. & Xie, X. S. Genome-wide study of mRNA degradation and transcript elongation in *Escherichia coli*. *Mol. Syst. Biol.* (2015).

29. Wang, M., Zhang, J., Xu, H. & Golding, I. Measuring transcription at a single gene copy reveals hidden drivers of bacterial individuality. *Nat. Microbiol.* **4**, 2118–2127 (2019).

30. Gillespie, D. T. Exact stochastic simulation of coupled chemical reactions. *J. Phys. Chem.* **81**, 2340–2361 (1977).

31. Meeussen, J. V. W. & Lenstra, T. L. Time will tell: comparing timescales to gain insight into transcriptional bursting. *Trends Genet. TIG* **40**, 160–174 (2024).

#### Supplemental Material

32. So, L. *et al.* General properties of transcriptional time series in *Escherichia coli*. *Nat. Genet.* **43**, 554–560 (2011).
33. Brewster, R. C. *et al.* The transcription factor titration effect dictates level of gene expression. *Cell* **156**, 1312–1323 (2014).
34. Zhang, Y., Ho, T. D., Buchler, N. E. & Gordân, R. Competition for DNA binding between paralogous transcription factors determines their genomic occupancy and regulatory functions. *Genome Res.* **31**, 1216–1229 (2021).
35. Cairns, B. R. The logic of chromatin architecture and remodelling at promoters. *Nature* **461**, 193–198 (2009).
36. Coulon, A., Chow, C. C., Singer, R. H. & Larson, D. R. Eukaryotic transcriptional dynamics: from single molecules to cell populations. *Nat. Rev. Genet.* **14**, 572–584 (2013).
37. Patange, S. *et al.* MYC amplifies gene expression through global changes in transcription factor dynamics. *Cell Rep.* **38**, 110292 (2022).
38. Yudkovsky, N., Ranish, J. A. & Hahn, S. A transcription reinitiation intermediate that is stabilized by activator. *Nature* **408**, 225–229 (2000).
39. Philips, S. J. *et al.* Allosteric transcriptional regulation via changes in the overall topology of the core promoter. *Science* **349**, 877–881 (2015).
40. Suter, D. M. *et al.* Mammalian genes are transcribed with widely different bursting kinetics. *Science* **332**, 472–474 (2011).
41. Zoller, B., Nicolas, D., Molina, N. & Naef, F. Structure of silent transcription intervals and noise characteristics of mammalian genes. *Mol. Syst. Biol.* (2015).
42. Harper, C. V. *et al.* Dynamic analysis of stochastic transcription cycles. *PLOS Biol.* **9**, e1000607 (2011).
43. Bremer, H. & Dennis, P. P. Modulation of chemical composition and other parameters of the cell at different exponential growth rates. *EcoSal Plus* (2008).
44. Henninger, J. E. *et al.* RNA-mediated feedback control of transcriptional condensates. *Cell* **184**, 207-225.e24 (2021).
45. Alon, U. *An Introduction to Systems Biology: Design Principles of Biological Circuits*. (Chapman and Hall/CRC, New York, 2019).

**Supplemental Material**

**TABLE S1. Contribution functions used in the study**

| Nascent RNA probes | Contribution function |
|---|---|
| For simulations/computational results | $g(\tau) = -\tau$ |
| *hb* CDS probes for data from[6,7] | $g(\tau) = \begin{cases} 1 & , -1 < \tau \le -\frac{3}{4} \\ \frac{-8\tau - 1}{5} & , -\frac{3}{4} < \tau \le -\frac{1}{8} \\ 0 & , -\frac{1}{8} < \tau \le 0 \end{cases}$ |
| *hb* P1 5'UTR probes for data from[7] | $g(\tau) = \begin{cases} 1 & , -1 < \tau \le -0.08 \\ \frac{-\tau}{0.08} & , -0.08 < \tau \le 0 \end{cases}$ |
| *hb* P2 3'UTR probes for data from[7] | $g(\tau) = \begin{cases} 1 & , -1 < \tau \le -0.99 \\ \frac{-\tau + 0.88}{0.11} & , -0.99 < \tau \le -0.88 \\ 0 & , -0.88 < \tau \le 0 \end{cases}$ |

**Supplemental Material**

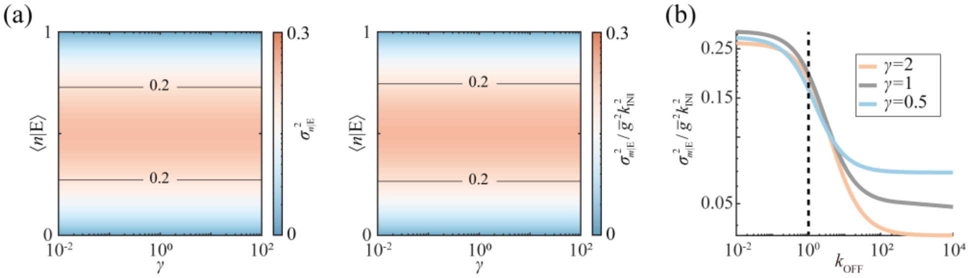

**FIG. S1. Intrinsic variance in coupled gene transcription.**

(a) Variance at the gene-state (left) and nascent-RNA (right) levels as functions of $\gamma$, and $\langle n \mid \mathrm{E} \rangle$, under a fixed environmental condition.

(b) Decay of nascent-RNA variance with $k_{\mathrm{OFF}}$ for representative $\gamma$ values ($\langle n \mid \mathrm{E} \rangle = 0.5$).

**Supplemental Material**

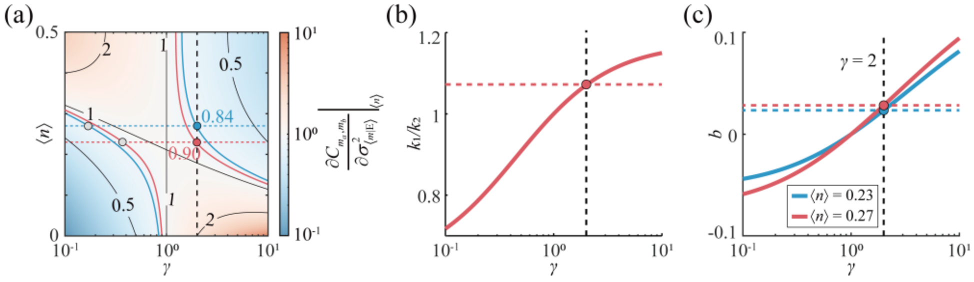

**FIG. S2. Methods for determining $\gamma$.**

(a) An alternative representation of Fig. 4(d). The partial slope of $\mathrm{Cov}(m_a, m_b)$ is plotted as a function of $\gamma$ and $\langle n \rangle$. Contours corresponding to experimentally measured partial slope values are overlaid, and their intersections with the respective $\langle n \rangle$ values are used to determine $\gamma$ for the system.

(b) The ratio of partial slopes of $\mathrm{Cov}(m_a, m_b)$ at two given $\langle n \rangle$ values as a monotonic function of $\gamma$. Comparing this curve with experimentally measured value uniquely determines $\gamma$ for the system.

(c) $f(\langle n \rangle, \gamma)$ at two given $\langle n \rangle$ values as monotonic functions of $\gamma$. For each curve, the intercept $b$ from the linear relationship between $\mathrm{Cov}(m_a, m_b)$ and $\sigma^2_{\langle m|\mathrm{E} \rangle}$ is used to uniquely determine $\gamma$ for the system.

#### Supplemental Material

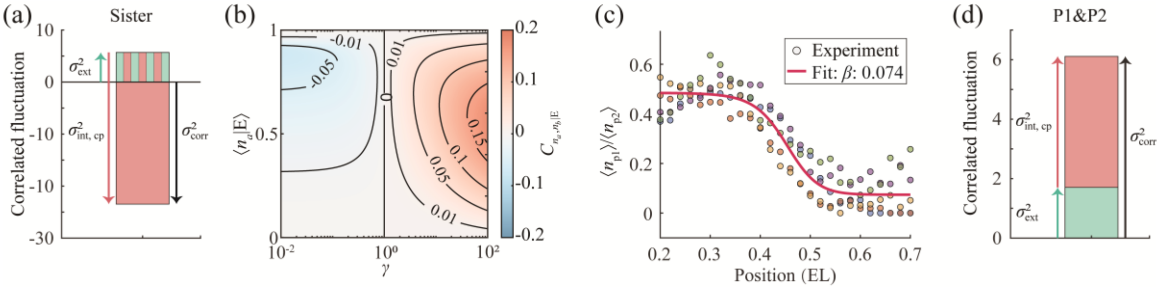

**FIG. S3. Noise decomposition for the *hb* gene.**

(a) Decomposition of correlated fluctuations for sister alleles (0.3−0.5 EL)

(b) Covariance between non-identical genes as a function of $\gamma$ and $\langle n_a \mid \mathrm{E} \rangle$ for $\beta = 0.1$.

(c) The ratio between $\langle n_{\mathrm{P1}} \rangle$ and $\langle n_{\mathrm{P2}} \rangle$ is plotted against the AP position. Data are fitted to a logistic function, with its minimum value used to estimate $\beta$. Dots of each color represent data from one of the five embryos at nc12–13.

(d) Decomposition of correlated fluctuations for P2 in comparison with P1 (0.3−0.5 EL)

#### Supplemental Material

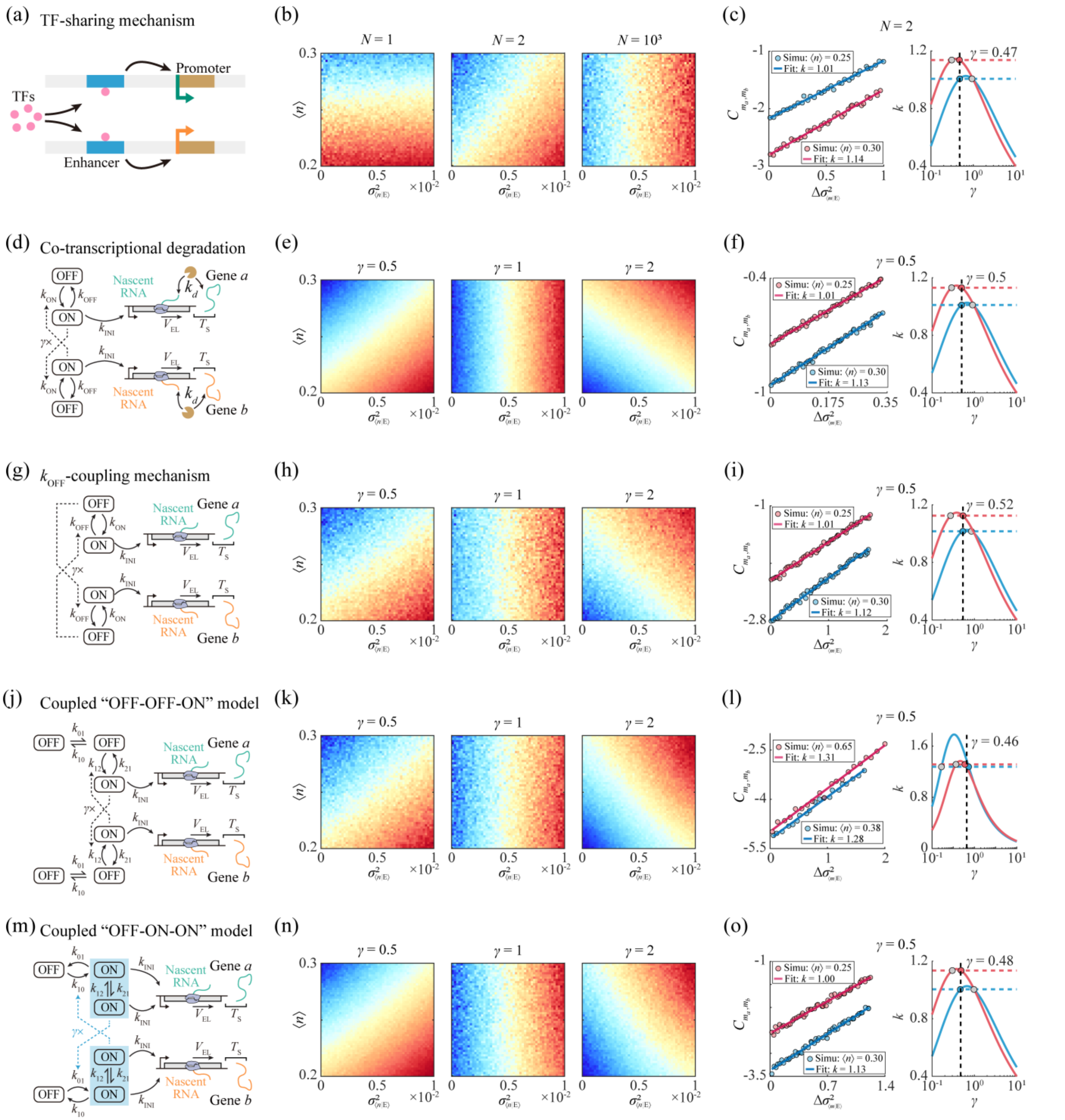

**FIG. S4. Verifying our noise decomposition framework for five alternative coupling mechanisms.**

(a, d, g, j, m) Schematics of different coupling mechanisms.

(b, e, h, k, n) Simulated total nascent-RNA covariance as a function of $\langle n\rangle$ and $\sigma^2_{\langle m|\mathrm{E}\rangle}$ across different coupling strength for each coupling mechanism.

(c, f, i, l, o) Estimation of $\gamma$ from the slopes of the linear relationships between total nascent-RNA covariance and $\sigma^2_{\langle m|\mathrm{E}\rangle}$ at given $\langle n\rangle$ values for each coupling mechanism.

#### Supplemental Material

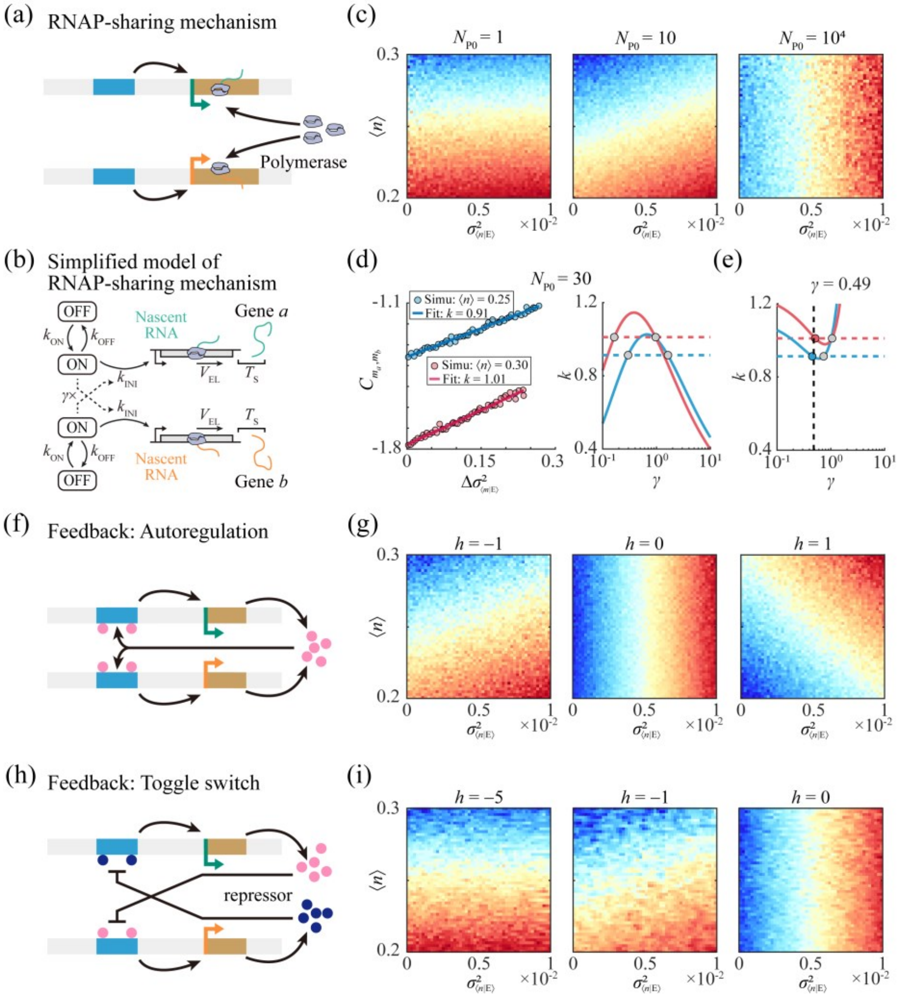

**FIG. S5. Verifying our noise decomposition framework for three additional coupling mechanisms.**

(a, f, h) Schematics of different coupling mechanisms.

(b) Schematic of a simplified model for RNAP-sharing mechanism.

(c, g, i) Simulated total nascent-RNA covariance as a function of $\langle n \rangle$ and $\sigma^2_{\langle m|\mathrm{E}\rangle}$ across different coupling strength for each coupling mechanism.

(d) Estimation of $\gamma$ from the slopes of the linear relationships between total nascent-RNA covariance and $\sigma^2_{\langle m|\mathrm{E}\rangle}$ for RNAP-sharing mechanism using $f$ from **Eq. (4)**. Results from different $\langle n \rangle$ values fail to yield a common solution for $\gamma$.

(e) Estimation of $\gamma$ from the slopes of the linear relationships between total nascent-RNA covariance and $\sigma^2_{\langle m|\mathrm{E}\rangle}$ for RNAP-sharing mechanism using $f$ estimated from the simplified model. Results from different $\langle n \rangle$ values successfully yield a common solution for $\gamma$.
